# Pan-cancer Drivers are Recurrent Transcriptional Regulatory Heterogeneities in Early-stage Luminal Breast Cancer

**DOI:** 10.1101/2020.03.30.017186

**Authors:** Shambhavi Singh, Matthew D. Sutcliffe, Kathy Repich, Kristen A. Atkins, Jennifer A. Harvey, Kevin A. Janes

**Affiliations:** Department of Biomedical Engineering, University of Virginia, Charlottesville, Virginia; Department of Radiology, University of Virginia, Charlottesville, Virginia; Department of Pathology, University of Virginia, Charlottesville, Virginia; Department of Imaging Sciences, University of Rochester Medical Center, Rochester, New York; Biochemistry & Molecular Genetics, University of Virginia, Charlottesville, Virginia

## Abstract

The heterogeneous composition of solid tumors is known to impact disease progression and response to therapy. Malignant cells coexist in different regulatory states that can be accessed transcriptomically by single-cell RNA sequencing, but these methods have many caveats related to sensitivity, noise, and sample handling. We revised a statistical fluctuation analysis called stochastic profiling to combine with 10-cell RNA sequencing, which was designed for laser-capture microdissection (LCM) and extended here for immuno-LCM. When applied to a cohort of late-onset, early-stage luminal breast cancers, the integrated approach identified thousands of candidate regulatory heterogeneities. Intersecting the candidates from different tumors yielded a relatively stable set of 710 recurrent heterogeneously expressed genes (RHEGs) that were significantly variable in >50% of patients. RHEGs were not strongly confounded by dissociation artifacts, cell cycle oscillations, or driving mutations for breast cancer. Rather, we detected RHEG enrichments for epithelial-to-mesenchymal transition genes and, unexpectedly, the latest pan-cancer assembly of driver genes across cancer types other than breast. Heterogeneous transcriptional regulation conceivably provides a faster, reversible mechanism for malignant cells to evaluate the effects of potential oncogenes or tumor suppressors on cancer hallmarks.

**Statement of significance:** Profiling intratumor heterogeneity of luminal breast carcinoma cells identifies a recurrent set of genes suggesting sporadic activation of pathways known to drive other types of cancer.

## Introduction

Approximately 80% of human tumors are epithelial carcinomas. Epithelial cells and progenitors proliferate considerably during normal tissue development, maintenance, and repair (1). Proper epithelial organization is enforced by basement membrane extracellular matrix (ECM), which becomes compromised during the epithelial cell-state changes underlying carcinomagenesis (2,3). Advanced carcinomas show considerable cell-to-cell variation in chromosomal gains–losses (4), overall mutational burden (5), and hybrid epithelial– mesenchymal traits (6). However, the state trajectory of carcinoma cells within large, rapidly progressing tumors is not stereotyped (7), complicating general interpretations of this variability. It is not known when intra-carcinoma cell heterogeneity meaningfully emerges, nor whether there might be common themes early in tumorigenesis that go on to diverge at later stages.

Thoroughly deconstructing intratumor heterogeneity requires transcriptomic approaches that can separate lineages and distinguish regulatory states with high cellular resolution (8,9). Conventional methods for single-cell RNA sequencing (scRNA-seq) either a) profile dozens– hundreds of cells at the maximum depth afforded by the single-cell sample, or b) shallowly scan (tens of) thousands of cells for molecular phenotyping (10). Regardless of the output, all leading approaches dissociate tumors into single-cell suspensions, requiring up to an hour of tissue processing and yielding a range of carcinoma proportions depending on cancer type (**Table 1**). The impact of these preparative steps on the transcriptomes of live cells is recognized (11) [and see also the accompanying study (12)], but they are considered to be an unavoidable tradeoff of the scRNA-seq approach.

**Table 1.**
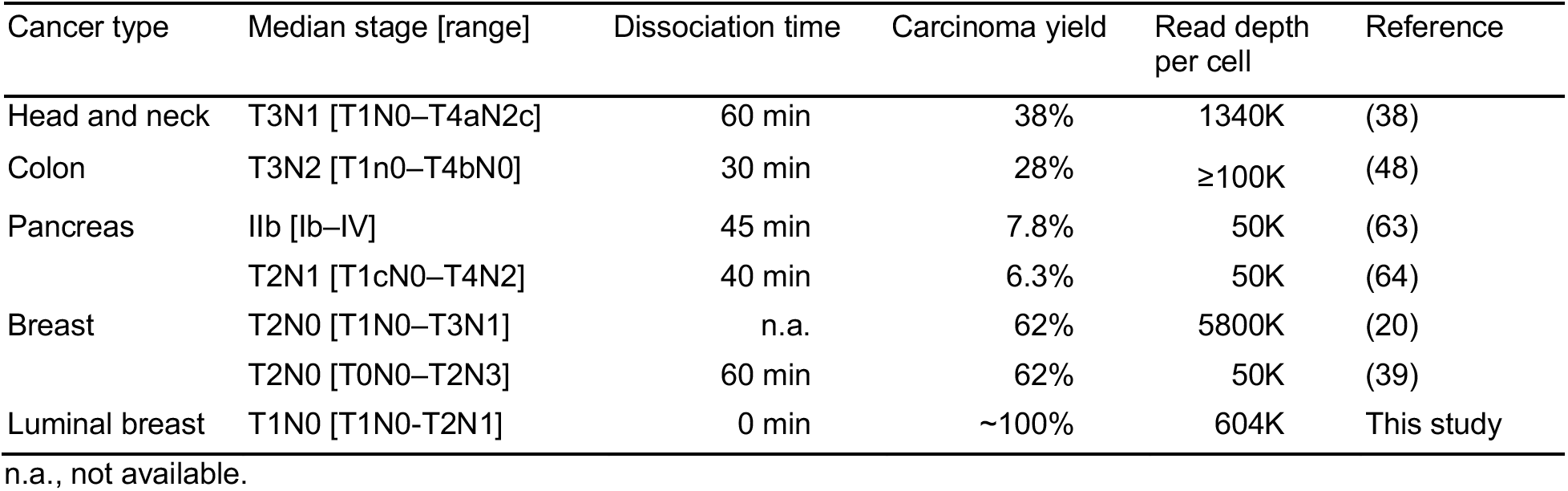
Transcriptomic studies of intra-carcinoma cell heterogeneity from primary clinical cases.

| Cancer type | Median stage [range] | Dissociation time | Carcinoma yield | Read depth per cell | Reference |
| --- | --- | --- | --- | --- | --- |
| Head and neck | T3N1 [T1N0–T4aN2c] | 60 min | 38% | 1340K | (38) |
| Colon | T3N2 [T1n0–T4bN0] | 30 min | 28% | ≥100K | (48) |
| Pancreas | IIb [Ib–IV] | 45 min | 7.8% | 50K | (63) |
|  | T2N1 [T1cN0–T4N2] | 40 min | 6.3% | 50K | (64) |
| Breast | T2N0 [T1N0–T3N1] | n.a. | 62% | 5800K | (20) |
|  | T2N0 [T0N0–T2N3] | 60 min | 62% | 50K | (39) |
| Luminal breast | T1N0 [T1N0–T2N1] | 0 min | ~100% | 604K | This study |

n.a., not available.

Considering the drawbacks, we (13) and others (14,15) have developed approaches orthogonal to scRNA-seq that combine high-sensitivity transcriptomics with laser-capture microdissection (LCM). Using LCM, histologically distinguishable cell types can be isolated with single-cell resolution from cryosections of snap-frozen tissue or tumor. Unfortunately, all LCM-based sequencing methods have substantially decreased sensitivity and technical reproducibility with fewer than 10 cells of microdissected material (14–16). To gain single-cell information, we reasoned that our 10-cell RNA sequencing (10cRNA-seq) method could productively identify carcinoma cell-regulatory heterogeneities if it were implemented as a type of fluctuation analysis called stochastic profiling (17). Previously, stochastic profiling was applied to 10-cell averaged gene-expression profiles collected repeatedly within a clonal cell line and measured by qPCR or microarray, leading to predicted single-cell heterogeneities that were subsequently validated (17–19). The distinct statistical properties of RNA sequencing data required design of a customized analytical pipeline for stochastic profiling by 10cRNA-seq.

Here, we combine 10cRNA-seq with an abundance-dependent overdispersion statistic that enables stochastic profiling of tumor cells *in situ.* Without any sample dissociation, we deeply profile carcinoma cell-to-cell heterogeneity in a cohort of five closely matched, late-onset and early-stage luminal breast cancers (**Table 1** and Supplementary Table ST1), obtaining data on 21,255 genes from 1400 carcinoma cells. The LCM component of 10cRNA-seq proved critical to obtain carcinoma heterogeneity profiles from cases with extensive immune infiltration. 10-cell pooling minimized the contribution of periodic transcripts that covary with cell-cycle phases. Stochastic profiling inferred 710 candidate transcripts that were recurrently heterogeneous in ≥50% of tumors. Among these RHEGs, 85% were retained when the same analysis was applied to three additional luminal breast cancers (20), suggesting bounds for reliable variability between similar tumors. The shared set of candidates was largely devoid of detachment-induced artifacts (21) and, surprisingly, breast-cancer driver genes (22). Recurrent heterogeneities were instead enriched in the collagen and matricellular constituents of a pancancer epithelial-to-mesenchymal transition (EMT) signature (23). The heterogeneities uncoupled canonical EMT marker genes that are usually linked at the population level. Most intriguing was an enrichment for dozens of non-cycling driver genes for cancers other than breast (24). Our results raise the possibility that early-stage luminal breast cancers experience a much broader landscape of oncogenes and tumor suppressors through transcriptional heterogeneity than indicated by the genomic lesions characteristic of the subtype.

## Materials and Methods

### Tissue procurement and processing

Human sample acquisition and experimental procedures were carried out in compliance with regulations and protocols approved by the IRB-HSR at the University of Virginia in accordance with the U.S. Common Rule. In accordance with IRB Protocol #19272, breast cancer samples were collected as ultrasound-guided core needle biopsies during diagnostic visits from participants who provided informed consent. Each core biopsy was cut into two pieces, freshly cryoembedded in NEG-50 medium (Richard-Allan Scientific) in a dry ice-isopentane bath, and stored at −80°C wrapped in aluminum foil. Cryosectioning and slide storage was performed exactly as described previously (13).

### Whole-exome sequencing

Genomic DNA was prepared from cryosectioned material by dissolving 8-μm sections in Buffer AL and purifying with the DNeasy Blood & Tissue kit (Qiagen). Whole-exome sequencing at 100x coverage was performed as a contract service with Genewiz. Raw BCL files were converted to fastq files with bcl2fastq v.2.19 and adapter trimmed with Trimmomatic v.0.38. Trimmed reads were mapped to the human reference genome (Build 37), and somatic variants were called using the Dragen Bio-IT Platform (Illumina) in somatic mode and a panel of normals to remove technical artifacts. Variants were removed if they were marked as common variants in dbSNP (Build 151) or if they had a non_cancer_AC > 5 in the gnomAD exome database v2.1.1. The filtered VCF was annotated with Ensembl Variant Effect Predictor (VEP) v95 for the Ensembl transcripts overlapping with the filtered variants. The set of consequence terms was defined by sequence ontology, and the term with the most severe impact was selected for each variant in downstream analysis. Summary of per-gene mutational annotations is available in Supplementary File S1, and summary of the per-mutation impact predictions and variant allele frequencies is available in Supplementary File S2.

### Rapid histology–immunofluorescence and laser-capture microdissection

For samples with low immune infiltration (UVABC1, UVABC2, and UVABC4) slides were stained and dehydrated as described previously (17). For samples with high immune infiltration (UVABC3 and UVABC5), slides were fixed immediately in 75% ethanol for 30 seconds, rehydrated quickly with PBS, and stained with a mixture of Alexa 488-conjugated KRT8 antibody (Abcam #ab192467, 1:20 dilution), YOPRO3 (Invitrogen #Y3607, 1:1000 dilution) and 1 U/ml RNAsin-Plus (Promega) in PBS for one minute. Slides were rinsed twice with PBS before dehydrating with 70% ethanol for 15 seconds, 95% ethanol for 15 seconds, and 100% ethanol for one minute and finally clearing with xylene for two minutes.

Slides were microdissected immediately on an Arcturus XT LCM instrument (Applied Biosystems) using Capsure HS caps (Arcturus). Cells were either visualized by brightfield microscopy (UVABC1, UVABC2, and UVABC4) or with a dual FITC/TRITC filter (UVABC3, UVABC5). The smallest spot size and typical instrument settings (~50 mW power and ~2 msec duration) yielded ~25 μm spot diameters capturing 1–3 breast carcinoma cells per laser shot.

For each biopsy, adjacent clusters of 10 cells were captured as 10-cell samples throughout multiple cryosections to access different regions of the tumor. In addition, a matched number of cells was captured nearby the 10-cell samples on the same LCM cap, extracted as a pool, and diluted into 10-cell equivalents that serve as measurement controls (pool-and-split controls). For each tumor, 10cRNA-seq datasets include 10-cell samples and matched pool-and-split controls captured across multiple days.

### RNA extraction and library preparation

RNA extraction and amplification of microdissected samples was performed as described previously (13). Briefly, RNA was eluted from the LCM caps by digesting with proteinase K, and oligo-dT primed cDNA was synthesized. Residual RNA was degraded by RNAse H (NEB) digestion, and cDNA was poly(A) tailed with terminal transferase (Roche). Poly(A)-cDNA was amplified using an AL1 primer (ATTGGATCCAGGCCGCTCTGGACAAAATATGAATTCTTTTTTTTTTTTTTTTTTTTTTTT) and a blend of Taq polymerase (NEB) and Phusion (NEB) for 30 cycles.

Poly (A) PCR-amplified samples were first assessed by quantitative PCR for exogenous ERCC spike in standards and endogenous genes (*GAPDH* and *RPL8* as loading controls and the epithelial marker *KRT8*) as previously described (13). New primers for this study were *KRT8* (Fwd: GCCGTGGTTGTGAAGAAGAT, Rev: CCCCAGGTAGTAAACTCCCC) and *RPL8* (Fwd: CCCAGCTCAACATTGGCAAT, Rev: ACGGGTCTTCTTGGTCTCAG). Samples were retained if the geometric mean of quantification cycles for the *GAPDH-RPL8* loading controls was within 3x the interquartile range of the median calculated across all 10-cell samples of the biopsy. Samples beyond that range were excluded because of over- or under-capture during LCM. For samples with increased immune infiltration, we additionally excluded samples with a detectable quantification cycle for the T cell marker *CD3D* (Fwd: TGCTTTGCTGGACATGAGACT, Rev: CAGGTTCACTTGTTCCGAGC).

Libraries for sequencing were re-amplified, purified, and tagmented as described previously (13). Briefly, each poly(A) PCR cDNA sample was re-amplified for a number of cycles where the amplification remained in the exponential phase (typically, 10 to 20 cycles). Reamplified cDNA was then twice purified with Ampure Agencourt XP SPRI beads. After bead purification, samples were quantified on a CFX96 real-time PCR instrument (Bio-Rad) using a Qubit BR Assay Kit (Thermo Fisher). Samples were diluted to 0.2 ng/μl and tagmented with the Nextera XT DNA Library Preparation Kit (Illumina).

### RNA sequencing

Libraries from 10-cell samples were multiplexed at an equimolar ratio, and 1.3 pM of the multiplexed pool was sequenced on a NextSeq 500 instrument with NextSeq 500/550 Mid/High Output v1/v2/v2.5 kits (Illumina) to obtain 75-bp paired-end reads. From the sequencing reads, adapters were trimmed using fastq-mcf in the EAutils package (version ea-utils.1.1.2-779), and with the following options: -q 10 -t 0.01 -k 0 (quality threshold 10, 0.01% occurrence frequency, no nucleotide skew causing cycle removal). Quality checks were performed using FastQC (version 0.11.8) and MultiQC (version 1.7). Data were aligned to the human transcriptome (GRCh38.84) along with reference sequences for ERCC spike-ins using RSEM (version 1.3.0) and Bowtie 2 (version 2.3.4.3). RSEM options for the 10cRNA-seq data also included the following: --single-cell-prior --paired-end. RSEM read counts were converted to transcripts per million (TPM) by dividing each value by the total read count for each sample and multiplying by 10^6^. Mitochondrial genes and ERCC spike-ins were not counted towards the total read count during TPM normalization.

### Molecular subtype assignments

Microarray-based PAM50 centroids and associated code were obtained from the UNC Microarray Database (https://genome.unc.edu/pubsup/breastGEO/PAM50.zip). To adapt the signature for RNA-seq data, RSEM aligned TPM data for TCGA breast tumors was obtained from the UCSC genome browser (25). Using a balanced number of estrogen receptor (ER)-negative and ER-positive tumors, median RNA-seq values for PAM50 genes were calculated and subtracted from the entire cohort for standardization. Standardized values were used to predict PAM50 subtypes by using downloaded centroids and code from the UNC Microarray Database (26). Successful model training was visualized by a principal component plot showing both training (microarray) and test (TCGA RNA-seq) data clustering by molecular subtype. The same median correction method was attempted for 10cRNA-seq data from UVABC tumors, but model calibration was unsuccessful due to a lack of ER-negative samples and some large differences in overall abundance of PAM50 genes between bulk and 10-cell data. As a substitute, 10cRNA-seq samples were scored for transcriptional modules associated with ESR1 (464 genes), ERBB2 (27 genes), and AURKA (229 genes) using the “subtype.cluster” function within the package “genefu” (version 2.16.0) in R. On the basis of these module scores, samples were subtyped as Luminal A (ESR1+, ERBB2–, AURKA–), Luminal B (ESR1 +, ERBB2–, AURKA+), HER2 (ERBB2+), and Basal (ESR1–, ERBB2–).

### UMAP projections

All UMAP projections were generated using the R package “umapr” (version 0.0.0.9001) with the following parameters: neighbors = 4, distance metric = “correlation”, minimum distance = 0. For UVABC embedding, RSEM counts for UVABC tumors were converted to TPM values and projected onto a UMAP using all endogenous transcripts. The same UMAP projection was re-colored by subtype classifications and batch number. Batch effects were excluded by visualizing TPM estimates of exogenous ERCC spike-in expression for all UVABC tumors on a separate UMAP. scRNA-seq data of breast tumors (20) were obtained as RSEM-aligned TPM values from the Gene Expression Omnibus (GSE75688). Epithelial carcinoma cells were separated from infiltrating cell types in the scRNA-seq data by selecting cells that expressed *KRT8* and *EPCAM* at TPM > 1. 10cRNA-seq UVABC samples and filtered scRNA-seq data (78 cells) were merged by transcript and projected on the same UMAP.

To account for tumor-specific differences in overall transcript abundance, we centered the expression of every transcript by the 25th quartile of its expression in 10-cell samples from each tumor. Quartile-centered samples for each tumor were then merged by transcript and projected onto a UMAP. From the merged quartile-centered data, expression of 710 RHEGs was extracted and projected onto a separate UMAP.

### Overdispersion-based stochastic profiling

10cRNA-seq analysis consisted of an identical algorithm applied separately to 28 10-cell samples and 20 pool-and-split controls from each UVABC tumor. RSEM values were rounded to integer counts, and transcripts with zero counts throughout were removed. Abundance-dependent expected expression and error models were generated separately for 10-cell samples and pool-and-split controls using the “knn.error.models” function in the package “scde” (version 1.99.4) with *k* nearest neighbors set to ¼ of the sample size for both sets. Only transcripts that had a minimum transcript count of 5 (min.count.threshold) in at least 5 samples (min.non.failed) were considered for model generation. From the abundance-adjusted error models, adjusted-variance estimates of overdispersion were calculated using the “pagoda.varnorm” function of the same package using the generated error models as input.

The variance was further adjusted to account for read-depth using the “pagoda.subtract.aspect” function. Transcripts with adjusted variances in 10-cell samples above the 5th percentile of adjusted variances in pool-and-split controls (the empirical type I error rate) were considered candidate heterogeneously expressed transcripts. Finally, transcripts with adjusted variances in the top 5th percentile of pool-and-split controls reflecting high measurement variability were filtered out of candidate lists. Overdispersion-based stochastic profiling was applied identically to each of the five UVABC cases to obtain candidate gene lists. Subsampled evaluations drew from the 28 10-cell observations without replacement. The computational assembly of 10-cell pools from scRNA-seq data (20) used the empirical type I error rate of UVABC4, which had the minimum adjusted variance of the cohort. Final candidate gene lists for every tumor are supplied along with RHEGs in Supplementary File S3.

### Cohort subsampling for RHEG estimation

The five UVABC tumors were exhaustively downsampled as groups of *n* = 1 (five total possibilities), 2 (10 total), 3 (10 total), 4 (five total), or 5 (one total) and intersected with the operational RHEG definition of transcript heterogeneities that recur in ≥50% of the cases considered: one for *n* = 1 or 2, two for *n* = 3 or 4, and three for *n* = 5.

### CNV inference

To predict CNVs from 10cRNA-seq, we used inferCNV, which corrects the input expression data for average gene expression based on normal reference cells and applies a moving average with a sliding window of 101 genes within each chromosome. Normal breast tissue gene-expression data was obtained from GTEx (27) as a reference dataset for copynumber variation. The reference GTEx data, UVABC 10cRNA-seq data, and a reference genome position file (GRCh38.86) were input to the “CreateInfercnvObject” function in the package “InferCNV” (version 1.0.3) in R (https://github.com/broadinstitute/inferCNV). The inferCNV object was then analyzed with the “infercnv::run” function with dynamic threshold denoising to infer copy-number variations as previously described (28).

### Inference of bulk differential expression

For a normal reference, single-cell RNA-seq data from three reduction mammoplasties (29) were downloaded and aligned by using RSEM identically as described above for 10cRNA-seq data. To generate pseudo-bulk normal profiles, gene-counts were summed across all individual cells from the same individual. For a bulk comparison with UVABC luminal breast tumors, gene counts were summed across pool-and-split controls measured by 10cRNA-seq and grouped by experimental day to generate biological replicates comprised of different regions of the same tumor. DESeq2 (version 1.24.0) was used to identify differentially expressed transcripts between pseudo-bulk normal data and tumor data using model design ~condition separately for each UVABC tumor. Genes with an adjusted *p* value less than 0.05 were considered significantly up-or downregulated between the two groups and are summarized in Supplementary File S4.

### Periodicity of cell-cycle RHEGs vs. non-RHEGs

We obtained 361 oscillating transcripts from Cyclebase 3.0 (30), reconciled aliases with official gene names, and intersected with the RHEGs from 10cRNA-seq, obtaining 24 shared transcripts. Next, using microarray data from synchronized HeLa cells (31), we identified probesets for 21 of the 24 shared transcripts along with those of the top 10 cycling transcripts according to Cyclebase 3.0 (30). The centered probeset data and HeLa pseudotime estimates are available through Cyclebase 3.0 for three complete cell cycles, but only the first two cycles show strong synchronization (30). We calculated the time interval above the mean value and compared it as a ratio to each of the adjacent time intervals below the mean value. For the ratio, the larger pseudotime interval (above or below the mean) was placed in the numerator. We calculated the skewness of the ratio distributions (indicating time-interval asymmetry above-vs.-below the mean) for the 21 RHEGs with identifiable probesets vs. the top-10 cycling transcripts and estimated uncertainty by bootstrapping.

### Monte Carlo simulations of three-state stochastic profiling

Monte Carlo simulations of stochastic profiling under the assumption of two regulatory states were performed in MATLAB with available software (16). The two-state model assumes a binomial distribution for the cellular regulatory-state dichotomy and log-normal distribution of measured transcripts. To build a three-state model reflecting cell cycle-regulated variation, we used a multinomial distribution to reflect cell-cycle fractions, two lognormal regulatory states for G1 and G2/M phases, and a uniform “S-phase” interval between the G1 and G2/M regulatory states. Two-or three-state distributions were compared against a null distribution of lognormal variation using the Kolmogorov-Smirnov test. The simulations for a parameter set were run 50 times to measure the median *p* values and the associated nonparametric confidence intervals. Stochastic sampling was deemed effective when the median *p* value for *F* ≠ 0 (multiple states) was less than 0.05 and the median *p* value for *F* = 0 (one state) was greater than 0.05. MATLAB code for the modified three-state distributions is available in Supplementary File S5.

### Gene signature overlaps with RHEGs

All overlaps were viewed using the “venn” function in the R package gplots (version 3.0.1.1), and intersections were obtained using the “intersect” function. Significance of overlap was calculated using the hypergeometric test in R through the “phyper” function and a total gene count of 20,000. For assessing detachment-induced artifacts in the RHEG gene set, we defined a detachment-induced gene signature as transcripts appearing at least twice among upregulated genes among four luminal breast-cancer lines grown in ultra-low attachment suspension (21). Breast driver genes were obtained from a larger gene list of cancer drivers (22) filtered for tissue of origin. We similarly assessed overlap with transcripts altered by CTCF knockout in luminal breast cancer cells (32), GATA3 target genes (33), EMT signature gene sets (23,34), and an aggregated pan-cancer driver gene set (22,35). For the last comparison, nine RHEGs were considered proximal to driver genes (Supplementary Table ST3) and treated as equivalent between the two sets. All mismatched gene aliases were corrected before overlap testing.

### Statistics

Sample sizes for stochastic profiling were determined by Monte Carlo simulation (16). Differences in genes detected per sample between 10cRNA-seq and scRNA-seq were assessed by Kolmogorov-Smirnov test using the “ks.test” function in R. Significance of overlaps between candidate genes identified in different UVABC tumors were assessed by Monte Carlo simulations that drew the total number transcripts per tumor randomly from a common pool of 14,824 genes (total transcripts eligible for overdispersion analysis in all tumors). Observed overlaps were compared with 1000 Monte Carlo simulations to estimate a *p* value, which was adjusted for multiple comparisons by using the Šidák correction. All overlaps between the RHEG set and other gene sets were assessed for significance by hypergeometric test using the “phyper” function in R and a background of 20,000 genes. Kolmogorov-Smirnov tests for Monte Carlo simulations for stochastic profiling were assessed using the “kstest” function in MATLAB. Hierarchical clustering was performed using “pheatmap” in R using Euclidean distance and “ward.D2” linkage.

### Data availability

10cRNA-seq from this study are available through the NCBI Gene Expression Omnibus (GSE147356, https://www.ncbi.nlm.nih.gov/geo/query/acc.cgi?acc=GSE147356 Reviewer token: wzcfouoijfodzij).

## Results

### Carcinoma-focused 10-cell profiling of early-stage luminal breast cancer

Women were selected for enrollment if they required ultrasound-guided biopsy for a suspected malignancy after screening mammography (BI-RADS 4C and higher). Just before diagnostic biopsy, we obtained written informed consent to collect an additional ultrasound-guided core sample, which was cryoembedded (13) within one minute of acquisition (**Fig. 1A** and **Table 1**). After clinical diagnosis of hormone-positive, *HER2-negative* breast cancer, we selected five cases that were as closely matched as possible (Supplementary Table ST1). The median tumor biopsy was Stage 1, Grade 3, and aged 63 years—the late-onset group for breast cancer. To define the mutational spectrum of each case, we performed whole-exome sequencing on DNA purified from the cryosectioned biopsy. The analysis identified 139–263 point mutations in 118–230 genes, with UVABC2 showing the highest burden and UVABC5 the lowest (Supplementary Files S1 and S2). Each case harbored mutations in multiple driver genes for luminal breast cancer (36). Albeit very focused, the cohort was both diverse and representative in terms of tumor genetics for the luminal subtype.

**Figure 1.**
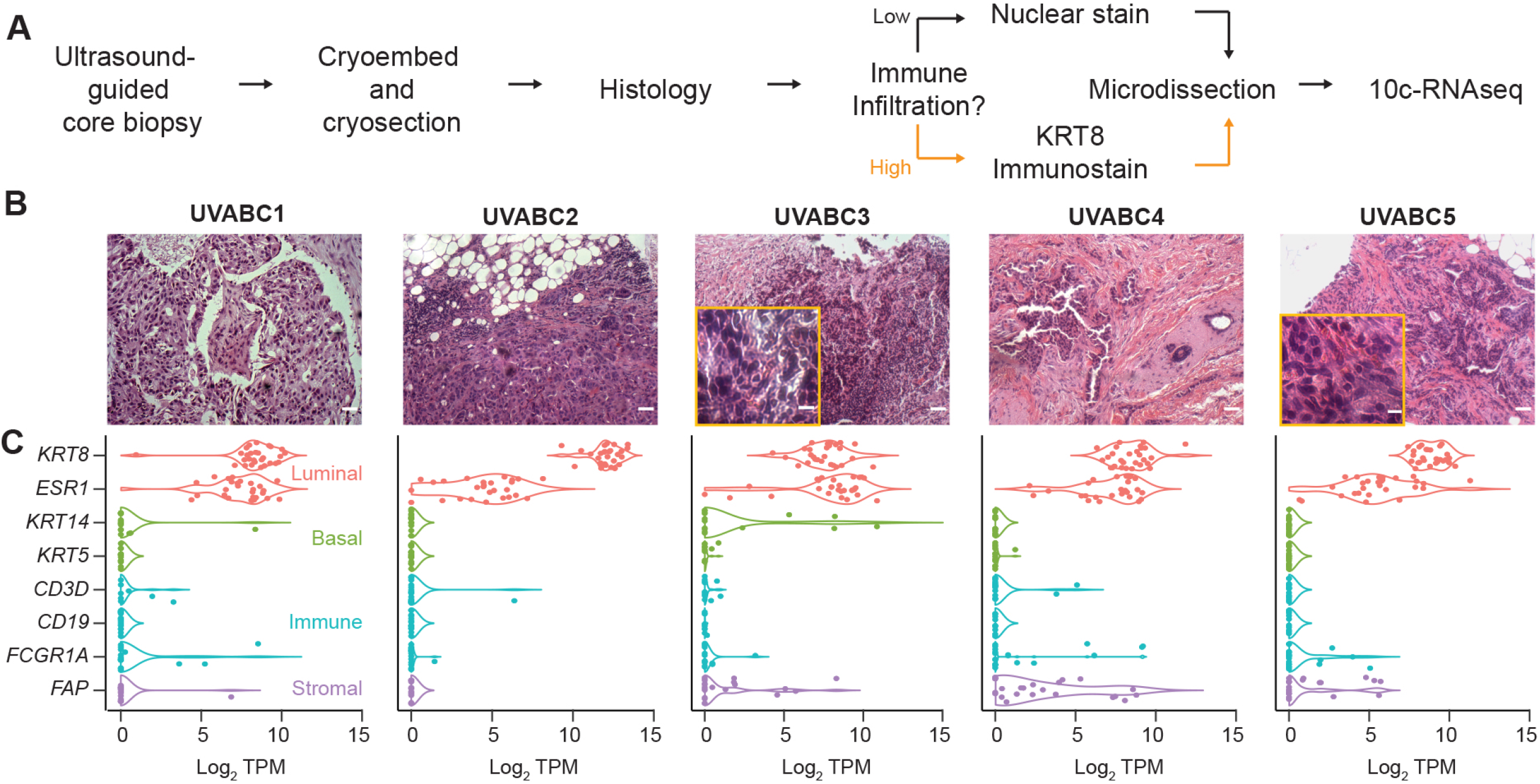
Focused transcriptional profiling of breast carcinoma cells without dissociation in early-stage tumor biopsies. **A,** Biopsy tissue was immediately cryoembedded and later cryosectioned for laser-capture microdissection. Tumor cells were visualized with either a rapid nuclear stain or KRT8 immunostain (Supplementary Fig. S1) before 10cRNA-seq (13). **B,** Tumor histology of the UVABC cohort visualized by hematoxylin-eosin staining. Two cases (marigold inset) showed increased tumor infiltrating lymphocytes requiring KRT8 immunostain-guided LCM. **C,** Selective capture of epithelial carcinoma cells assessed by marker transcripts for luminal cells (*KRT8, ESR1*), basal cell (*KRT14, KRT5*), immune cells (*CD3D, CD19, FCGR1A*), and stromal content (*FAP*). Scale bar is 80 μm (**B**) or 10 μm (**B**, inset).

Despite histologic similarities in gross tumor characteristics, we noted elevated lymphocyte infiltration in two cases (UVABC3 and UVABC5) that rendered them problematic to microdissect by nuclear histology alone (**Fig. 1B** and Supplementary Table ST1). Therefore, we devised an immuno-LCM procedure that combines an Alexa Fluor 488-conjugated, high-affinity monoclonal antibody against KRT8 with the red-orange nucleic acid stain YOPRO3 (see Materials and Methods). A one-minute incubation with the antibody-dye cocktail was sufficient to resolve KRT8-positive carcinoma cells from KRT8-negative stromal cells and the YOPRO3-negative autofluorescence of tissue collagen (Supplementary Fig. S1A). We could not find any evidence that the antibody or dye interfered with the critical early steps of 10cRNA-seq (Supplementary Fig. S1B) (13).

For each case, we collected 28 random pools of 10 carcinoma cells located throughout cryosections of the core biopsy. Pools were assembled as local 10-cell groups that reflect both clonal and microenvironmental heterogeneity. We recorded the spatial position of all cells microdissected in the 10-cell samples to leave open the possibility of retroactively linking transcriptomic changes to histological or topological features of the tumor. Samples were deeply sequenced at 6.04 ± 0.75 million reads per 10-cell pool to ensure saturation of gene detection and provide maximum sensitivity for identifying non-carcinoma contaminants. Across all cases, we found that the luminal markers *KRT8* and *ESR1* predominated by transcripts per million (TPM), whereas markers for myoepithelial cells (*KRT14, KRT5*), T cells (*CD3D*), B cells (*CD19*), and macrophages (*FGCR1A)* were rarely detected (**Fig. 1C**). Even though desmoplasia was marked for all but one case (Supplementary Table ST1), fibroblast-like markers were detectable in only ~14% of 10-cell samples (using log2 *FAP* > 5 transcripts per million [TPM] ≈ 0.8 copies per cell (13) as a stringent threshold). Overall, the observations confirmed the fidelity of (immuno-)LCM for isolating spatially-resolved, carcinoma-specific transcriptomic profiles with minimal disruption of tumor architecture.

### 10cRNA-seq transcriptomes retain the inter-tumor and intra-tumor heterogeneity of luminal breast carcinoma cells profiled by scRNA-seq

As a first assessment, the 10cRNA-seq data were visualized by uniform manifold approximation and projection (UMAP) (37). Consistent with past descriptions of carcinoma heterogeneity by scRNA-seq (20,38,39), we found that 10cRNA-seq data clustered tightly by patient (**Fig. 2A**). Batch effects were not evident within the patient clusters (Supplementary Fig. S2A–E). Furthermore, we did not observe any clustering in a separate UMAP visualization using only the data from ERCC spike-ins added to every sample at the time of RNA extraction (Supplementary Fig. S2F). The observed separation of 10cRNA-seq transcriptomes (**Fig. 2A**) thus reflects bona fide inter-tumor differences between cases.

**Figure 2.**
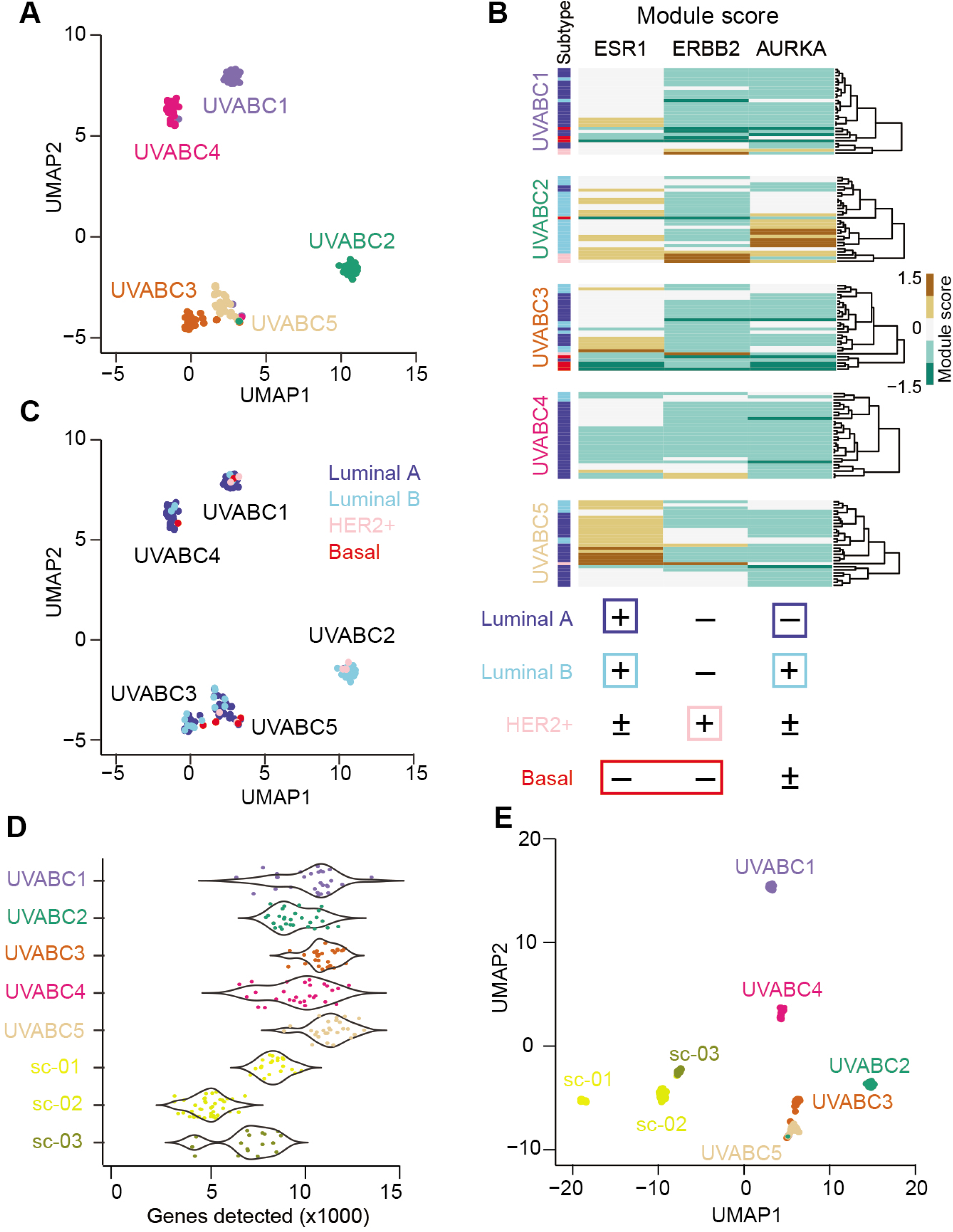
10-cell transcriptomes of luminal breast carcinomas are heterogeneous among and within tumors. **A,** UMAP embedding of 10cRNA-seq samples from the UVABC cohort colored by tumor. **B,** A three-signature classification system for identifying molecular subtypes of breast cancer in 10cRNA-seq data. Module scores were used to classify the subtype of each sample as indicated. **C,** Molecular subtype classifications of the 10cRNA-seq samples projected as in **A. D,** Genes detected by 10cRNA-seq in the UVABC cohort compared to three luminal tumors profiled by scRNA-seq (sc-01, sc-02, sc-03) (20). *p* < 10^-15^ by K-S test. **E,** UMAP embedding of tumors profiled by 10cRNA-seq and scRNA-seq.

Next, we sought to classify individual 10cRNA-seq samples into intrinsic breast cancer subtypes. We adapted the microarray-based PAM50 classification of subtypes (26) to RNA-seq (25), but the 50-gene signature was not robust enough for 10cRNA-seq observations (Supplementary Fig. S3). A similar distinction between bulk and 10-cell projections was noted in the accompanying study (40). As a substitute, we used expression-signature modules (41) associated with *ESR1, ERBB2,* and *AURKA* as proxies for the hormone, *HER2,* and proliferative status of each 10-cell sample (see Materials and Methods). Within patients, there was considerable variability in module scores (**Fig. 2B**), corroborating an earlier scRNA-seq study of multiple breast-cancer subtypes (20). Although most 10-cell samples were classified as luminal A or luminal B subtype, all cases but UVABC4 contained observations scoring more strongly to other subtypes (**Fig. 2C**). Three cases (UVABC1–3) harbored instances of all four subtypes, analogous to scRNA-seq observations in glioblastoma (28). The clustering of mixed classifications implied that other patient-specific gene programs were dominant in the UMAP organization. Variations in subtype class were repeatedly documented in nearby samples microdissected from the same histologic section (Supplementary Fig. S4). Even for early-stage breast tumors, the 10cRNA-seq data suggested that local variations in regulatory state are pervasive.

Toward a more-direct comparison of 10-cell data with single-cell measurements of gene expression, we extracted 78 *KRT8^+^EPCAM^+^* carcinoma cells from three cases of luminal breast cancer profiled by scRNA-seq (20) (see Materials and Methods). As previously reported by us and others (13,14,42,43), there were significantly more transcripts detected in local 10-cell pools compared to singly isolated cells (10,066 ± 1,416 genes vs. 5,957 ± 1,824 genes; *p* < 10^-15^ by K-S test; **Fig. 2D**). Overall, 3319 transcripts found in 10cRNA-seq pools were entirely undetected by scRNA-seq. Notwithstanding the differences in gene coverage, when scRNA-seq and 10cRNA-seq samples were projected on a shared UMAP, the separation between methods was comparable to that among patients (see Materials and Methods; **Fig. 2E**). Together, the data argue that 10-cell pooling does not dilute out the cell-to-cell and tumor-to-tumor heterogeneities recognized by scRNA-seq.

### Stochastic profiling by 10cRNA-seq identifies candidate regulatory heterogeneities

To go beyond qualitative descriptions of molecular subtype and inter-tumor differences, we sought to adapt the theory of stochastic profiling (16,17) to 10cRNA-seq (13). Stochastic profiling detects single-cell regulatory heterogeneities by statistically analyzing measurement fluctuations among small pools of cells collected randomly within a cell type (17). Samples with different proportions of cells in high-vs.-low regulatory states from a ~10-cell pool will create skewed distributions that reflect characteristics of the underlying single-cell population (19). Compared to earlier microarray-based studies, stochastic-profiling fluctuation analysis using RNA-seq brings new opportunities and challenges. One advantage is that RNA-seq is not biased by the position or quality of hybridization probes [as discussed in (13)]. However, transcript estimates are sensitive to read depth, and low-abundance genes are susceptible to noise from counting statistics. Rare transcripts are a key hurdle, because sample-to-sample fluctuations will be partially obscured by the detection limit, which combines instances that are “absent” with those that are “not counted “ (**Fig. 3A**). We surmounted these hurdles by extracting a module for estimating “abundance-dependent dispersion” from the SCDE package (44,45) and redeploying it as a separate inference tool for stochastic profiling. The module relates the squared coefficient of variation (CV) among replicates of each gene in a study (here, a patient) to the abundance magnitude of that gene, building a model for the within-study variance expected of a transcript at a given abundance (**Fig. 3B**). The variance of each transcript is then normalized by the expected variance for that transcript’s abundance, yielding an adjusted variance (Varadj) that acts as an overdispersion score. For high-abundance genes, overdispersed transcripts such as *MYL12B* show multiple modes or heavier tails than expected (compare with *GABARAP,* **Fig. 3C** and **3D**). Low-abundance genes with overdispersion such as *TXNRD3* are skewed by multiple instances of moderate-to-high TPM (compare with *DSCR3*, **Fig. 3E** and **3F**). The dispersion module incorporates discrete random variables (negative binomial and Poisson) to model aligned reads and dropouts. The Var_adj_ overdispersion score thus provides a principled metric for stochastic-profiling analysis of 10cRNA-seq data.

**Figure 3.**
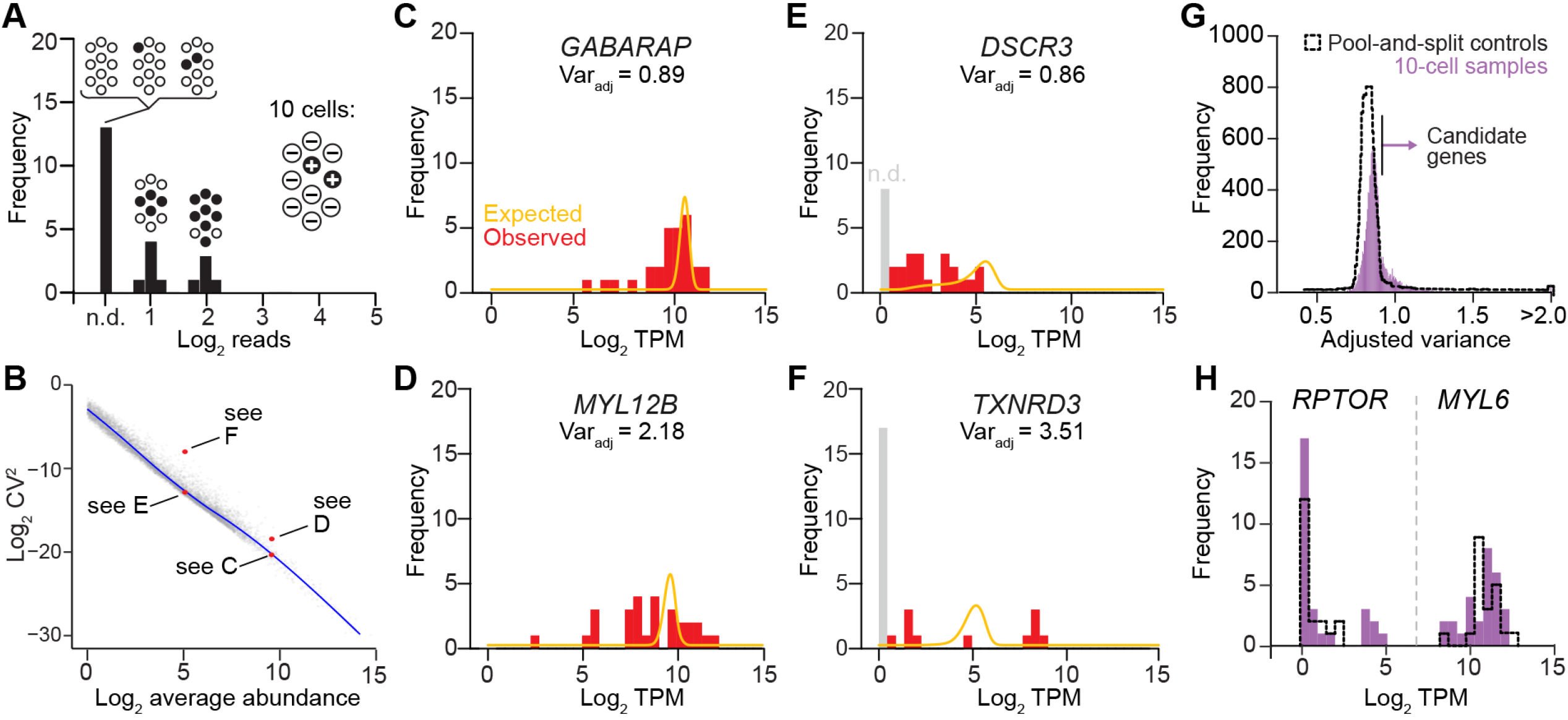
Stochastic profiling by 10cRNA-seq through abundance-dependent overdispersion statistics. **A,** Illustration of theoretical left-censored transcript, which is heterogeneously expressed at one copy per cell (filled, +) and measured with 50% efficiency. Low-frequency mixtures will be equivalently not detected (n.d.). **B,** Transcript dispersion is inversely related to abundance. Example transcripts with similar abundance but different dispersion (red) are annotated by the **Fig. 3** subpanel in which they appear. The blue trace indicates a smoothing cubic spline fit of the summarized 10cRNA-seq fluctuations per transcript (gray) that relates the coefficient of variation (CV) of each gene to its average abundance. **C** and **D,** 10cRNA-seq sampling fluctuations for high-abundance transcripts with expected dispersion (*GABARAP*, **C**) and overdispersion (*MYL12B,* **D**). **E** and **F,** 10cRNA-seq sampling fluctuations for low-abundance transcripts with expected dispersion (*DSCR3,* **E**) and overdispersion (*TXNRD3,* **F**). **G,** Distribution of adjusted variance scores for each gene measured transcriptomically in separate 10-cell samples (purple) compared to pool-and-split controls estimating technical variation (black dashed). The arrow indicates the upper 5th percentile of adjusted variance for the pool-and-split controls, which is used as the 10-cell type I error rate for identifying candidate genes. **H,** Examples of low-abundance (*RPTOR)* and high-abundance (*MYL6)* transcripts deemed to be significantly overdispersed given their relative abundance in 10-cell samples (purple) and their technical variation in pool-and-split controls (black dashed). For **C–F**, continuous traces (marigold) indicate the idealized dispersion expected given the observed transcript abundance, and the adjusted variance (Varadj) measure of dispersion is reported. Gray bars represent dropout events, which are not detected (n.d.) and modeled by a separate posterior (44,45). For **B–H**, data from UVABC4 were used as representative examples.

In conventional scRNA-seq, each cell is considered as an N-of-1 observation that convolves biological variability and technical noise. By contrast, stochastic profiling quantifies technical noise explicitly by pool-and-split analysis of 10-cell equivalents from hundreds of carcinoma cells microdissected in the same vicinity as the samples. We sequenced 20 pool-and-split controls in parallel with the 28 10-cell samples, analyzing the controls separately to construct a null distribution for transcript overdispersion in each tumor. The upper 5th percentile in the null model was used as an empirical type I error rate that defined the critical overdispersion cutoff for 10-cell samples—transcripts above this threshold in the 10-cell samples (but not in the null) were considered candidate regulatory heterogeneities (**Fig. 3G**, **3H**, and Supplementary Fig. S5).

### Stochastic profiling identifies recurrent transcriptional regulatory heterogeneities

By abundance-dependent dispersion, stochastic profiling identified 9206 candidate heterogeneities in the UVABC cohort, 161 transcripts of which were undetected by scRNA-seq (20) (Supplementary File S3). The extent of 10-cell sampling was more than sufficient to define the prevalent regulatory heterogeneities in each tumor—when datasets were subsampled, the recall of UVABC candidates saturated when 20 or greater 10-cell samples were used (Supplementary Fig. S6). We considered the contribution of heritable differences among tumor subclones in two ways, inferring copy-number variations from either 10cRNA-seq observations or bulk whole-exome sequencing (Supplementary Fig. S7 and S8). There was good agreement between the two approaches, including the inferred loss of 8p (all) and 11 q (UVABC1), along with gains in 1q (UVABC1,2,4) and 8q (UVABC2–4). Neither inference indicated that candidate heterogeneities were biased toward amplified or deleted loci. However, UVABC1—the case with the most-extensive chromosomal instability—was suggestive (*padj* = 0.07), supporting the study decision to avoid advanced tumors. For early-stage cancer, stochastic profiling by 10cRNA-seq nominates transcriptional regulatory heterogeneities that not appreciably biased by undersampling or confounded by subclonal genomic variation.

We next asked whether candidate heterogeneities showed any bias toward transcript upregulation or downregulation. To estimate bulk tumor transcriptomes for the UVABC cohort, we aggregated 10cRNA-seq pool-and-split controls from different days and spatially separated cryosections (see Materials and Methods). Normal reference samples were compiled bioinformatically using plate-based scRNA-seq of breast epithelia from three reduction mammoplasties (29). Differential expression analysis identified 5912–7876 transcripts with bulk abundance differences compared to the normal reference (Supplementary File S4). There were more downregulated transcripts by this comparison, but stochastic-profiling candidates were clearly enriched among those that were upregulated (Supplementary Table ST2). The bias is expected given that a regulatory burst of transcription will affect cellular mRNA abundances more rapidly than transcriptional silencing. Importantly, 58–67% of candidates showed no detectable difference in average abundance, illustrating that most regulatory heterogeneity escapes bulk differential expression, as further corroborated by the accompanying study (40).

Overall, 3627 candidates by stochastic profiling were exclusive to one of five patients, laying bare the extraordinary challenge of interpreting malignant cell-state heterogeneity beyond well-known markers of differentiation (9). Encouragingly, when the candidates were intersected, we noted a significant enrichment of transcripts that appeared repeatedly in three or more breast-cancer cases (**Fig. 4A**). Such transcripts were classified as recurrent heterogeneously expressed genes (RHEGs), for which there were 710 in total. Generalizing the RHEG definition to candidates appearing in >50% of the cases considered, we examined the stability of RHEG numbers by subsampling the UVABC cases (see Materials and Methods). As the quantity of patients increased, RHEGs stabilized in the range of 500–1500 for this highly circumscribed cohort (**Fig. 4B** and Supplementary Table ST1). We examined the generality of RHEGs by building simulated datasets of 10 carcinoma cells from scRNA-seq of three luminal breast cancers described earlier (20) (Supplementary Fig. S9A). Among the 610 RHEGs that were detectable by scRNA-seq, 85% were retained as RHEGs when the three additional cases of luminal breast cancer were considered (Supplementary Fig. S9B). RHEGs thus provide a robust conceptual framework for prioritizing cell-state regulatory heterogeneities identified in vivo (12,40).

**Figure 4.**
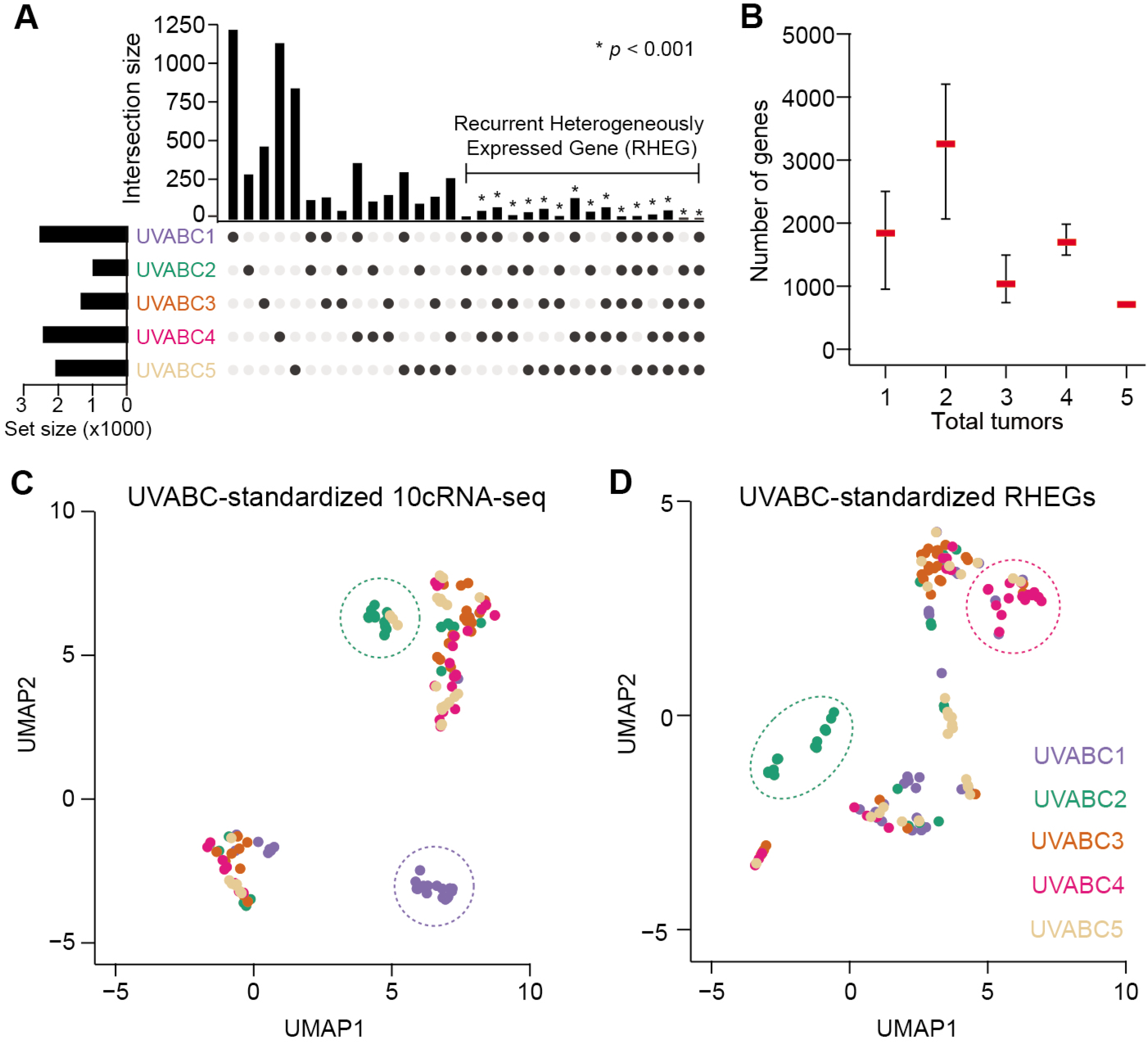
Intersecting candidate regulatory heterogeneities across tumors yields a set of recurrent heterogeneously expressed genes (RHEGs). **A,** Gene overlaps between and among cases in the UVABC cohort. Transcripts belonging to each tumor are available in Supplementary File S1. Enriched categories were assessed statistically by Monte-Carlo simulation (see Materials and Methods). **B,** RHEG size as a function of the number of UVABC tumors included. RHEGs were defined as transcripts that appear in >50% of the tumors included. Data are shown as the median ± range of _5_C_*n*_ combinations of *n* tumors in the UVABC cohort. **C** and **D,** UMAP embedding of patient-standardized 10cRNA-seq transcriptomes (**C**) or RHEGs (**D**). Patient-enriched clusters are highlighted with dashed ovals.

To evaluate RHEGs as an organizing principle, we revisited the UMAP visualization of UVABC cases from the standpoint of regulatory heterogeneity (**Fig. 2A**). Because abundance-dependent dispersion evaluates fluctuations local to each tumor (**Fig. 3**), it was first necessary to standardize the 10cRNA-seq transcriptomes separately and regenerate the UMAP (see Materials and Methods). Tumor-specific standardization intermingled the 10cRNA-seq observations considerably, but two clusters remained enriched in samples from UVABC1 and UVABC2 (**Fig. 4C**). When the same approach was applied using RHEGs exclusively, we observed a projection that was different from when the whole 10cRNA-seq transcriptome was used (**Fig. 4D**). For the same UMAP parameters, samples were more distributed than clustered, with the UVABC1-enriched cluster disappearing and a new UVABC4-enriched cluster appearing. This analysis suggested that RHEGs could be used as a lens to refocus transcriptome-wide heterogeneity on the most-robust variations.

### RHEGs are not dominated by cell-cycle covariates

In scRNA-seq, the most-overarching contributor to heterogeneity is the phase of the cell cycle (38,46). However, it was not obvious whether such single-cell variations would also overrepresent in RHEGs derived from 10-cell pools. Using a panel of 863 transcripts associated with replicating cells (46), we identified 62 among the 710 RHEGs, a significant overlap (*p* < 10^-6^ by hypergeometric test) but one comprising <10% of the list overall. Most of the overlapping genes were expressed acutely during one cell-cycle transition (e.g., G1/S, G2/M), which is akin to the two-state expression models foundational for the theory of stochastic profiling (16,17,19). When the cell-cycle search was restricted to 361 oscillating transcripts (30), the RHEG intersection reduced to 24 genes (*p* < 0.01 by hypergeometric test). Moreover, when we compared the periodicity of the overlapping genes to the most-symmetrically cycling transcripts, we found that RHEGs were significantly more imbalanced in their oscillations, implying bias toward one phase of the cell cycle (see Materials and Methods; Supplementary Fig. S10A and S10B and Supplementary File S6). The lack of oscillators directly proportional to the cell cycle is consistent with Monte-Carlo simulations of stochastic profiling that model a three-state distribution corresponding to G1, S, and G2/M populations (Supplementary Fig. S10C–F and Supplementary File S5). Stochastic-profiling theory thus bolstered our conclusion that genes tracking with cell-cycle oscillations are filtered out when transcriptomes are measured as 10-cell pools.

### RHEGs are largely devoid of detachment artifacts and influence from breast-cancer driver genes

We also considered other trivial explanations for genes in the RHEG set. Although tumors were not dissociated, it was possible that detachment-like regulatory variation was induced locally and rapidly from the tissue damage of the biopsy procedure. Using a 547-gene signature of suspension culture in luminal breast cancer cells (21), we intersected with the RHEG set and found that only 14 were shared (**Fig. 5A**). Thus, our clinical-procurement and sample-handling procedures had avoided detachment-like damage responses in the breast carcinoma cells.

**Figure 5.**
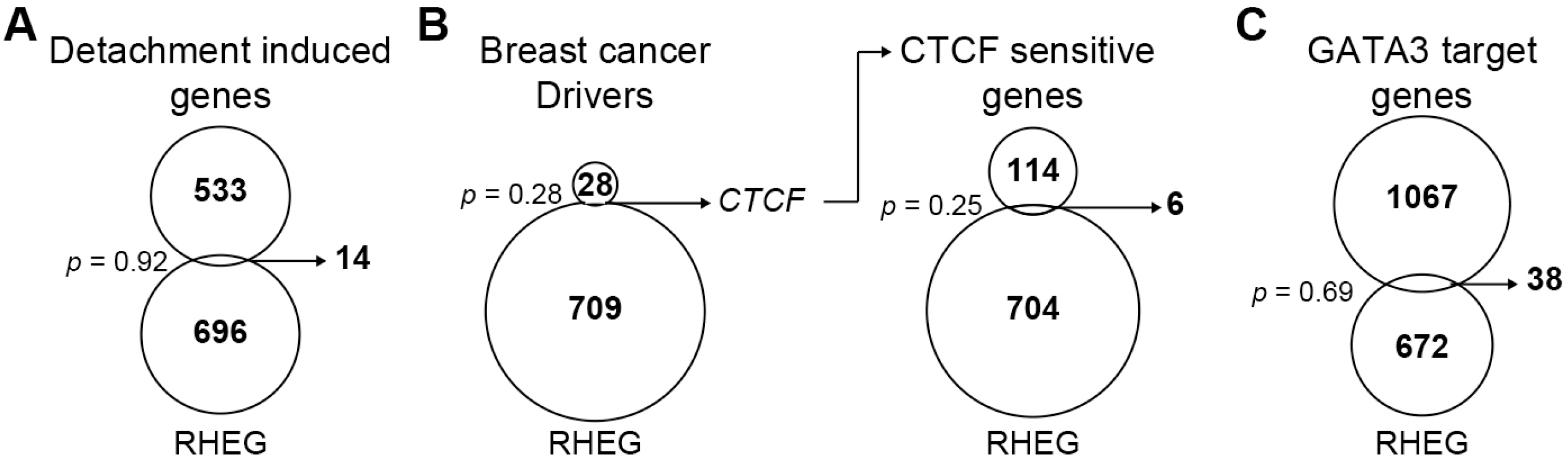
RHEGs have little in common with detachment signatures or mutational drivers of breast cancer. **A–C,** Venn diagrams intersecting the UVABC RHEG set with a cell-detachment signature (21) (**A**), a set of robust drivers for breast cancer (22) (**B**, left), a list of transcripts altered by CTCF knockout in a luminal breast cancer cell line (32) (**B**, right), and a list of GATA3 target genes (33) (**C**). Statistical significance of overlaps was assessed by the hypergeometric test.

Metascape (47) analysis revealed a significant overlap between the RHEG set and genes associated with luminal breast cancer (*q* < 10^-4^; Supplementary File S7). Therefore, we looked at known breast cancer drivers with the premise that mutations may arise subclonally in a breast carcinoma and disrupt abundance of the encoded transcript. Among 29 robust driver genes for breast cancer (22), only one was shared with the RHEG set: *CTCF* (**Fig. 5B**, left). As an insulator protein, CTCF abundance changes could cause secondary transcriptional alterations; however, we did not observe any enrichment for conserved CTCF-sensitive genes (32) in the RHEG set (**Fig. 5B**, right). The same was true when we separated candidate heterogeneities for UVABC2, which has an F416L mutation in CTCF at an allele fraction of 22% (Supplementary Fig. S11A and Supplementary File S2). Although not classified as a RHEG, we also investigated transcriptional targets for the most-prevalent transcription factor mutated in luminal breast cancer, GATA3 (22). We again found no RHEG enrichment among transcripts altered by mutant GATA3 in luminal breast cancer cells (33) (**Fig. 5C**). However, when candidates were separated for UVABC4, which has a D48Y mutation in GATA3 at an allele fraction of 23%, there was a detectable overlap (Supplementary Fig. S11B and Supplementary File S2). Stochastic profiling can thus detect the consequences of fractionally mutated transcription factors in primary cancers—the lack of association with RHEGs supported that RHEGs were more than a reflection of known sources of cell-to-cell heterogeneity in cancer.

### RHEGs are enriched for EMT signatures and cofluctuate with canonical EMT markers

scRNA-seq of dissociated tumors has identified (partial-)EMT states in some carcinomas (38,39) but not others (48). For breast cancer, changes along the EMT spectrum are mostly described in hormone-negative cell lines, but more-recent work reports EMT-like activation patterns in 65–85% of primary luminal breast cancers (49). We intersected a pan-cancer EMT signature (34) with the RHEG set and found significant overlap in multiple collagens, matricellular proteins, and other transcripts in the signature (**Fig. 6A**). The data suggest that ECM dysregulation in these tumors is jointly mediated by the carcinoma cells together with cancer-associated fibroblasts. RHEG enrichment was also found with an independently derived EMT signature (23) (Supplementary Fig. S12A), reinforcing the result. Metascape (47) analysis additionally supported a modest enrichment in target genes for the EMT effector, SNAI1 (*q* < 0.05; Supplementary File S7). Formally, none of the canonical EMT regulators [*ZEB1* (M), *ZEB2* (M)] or markers [*CDH1* (epithelial, E), *VIM* (mesenchymal, M), *FN1* (M)] were RHEGs, even though *VIM* was a candidate heterogeneity in two cases (UVABC3, UVABC5) and *FN1* a candidate in one case (UVABC3). We clustered these canonical EMT transcripts with the EMT RHEGs and found clear separation of E- and M-associated transcripts among 10-cell pools along with several notable subclusters by gene and by sample (**Fig. 6B**). The organization by patient was unexpected; for example, UVABC2 showed the most evidence for the E state, even though it was one of the most-advanced stage tumors of the cohort (Supplementary Table ST1). Reciprocally, 10-cell profiles of the high-grade UVABC3 tumor were no more scattershot in EMT transcripts than UVABC4 (grade 2) or UVABC5 (grade 1) (**Fig. 6B**). Among samples with M characteristics, *ZEB2* appeared prominently among samples abundant for some transcripts (*FN1, COL6A1, SPARC, VIM)* but not others (*COL5A2,* TAGLN). M-state fragmentation was also observed in UVABC1, which was predominated by samples positive for *VIM, SPARC,* and *SERPING1* but negative for CTSK, TAGLN, and mesenchymal collagens (**Fig. 6B** and Supplementary Fig. S12B). The RHEG set thereby provided a transcriptomic resource for posing questions about EMT regulatory patterns in early-stage luminal breast cancers.

**Figure 6.**
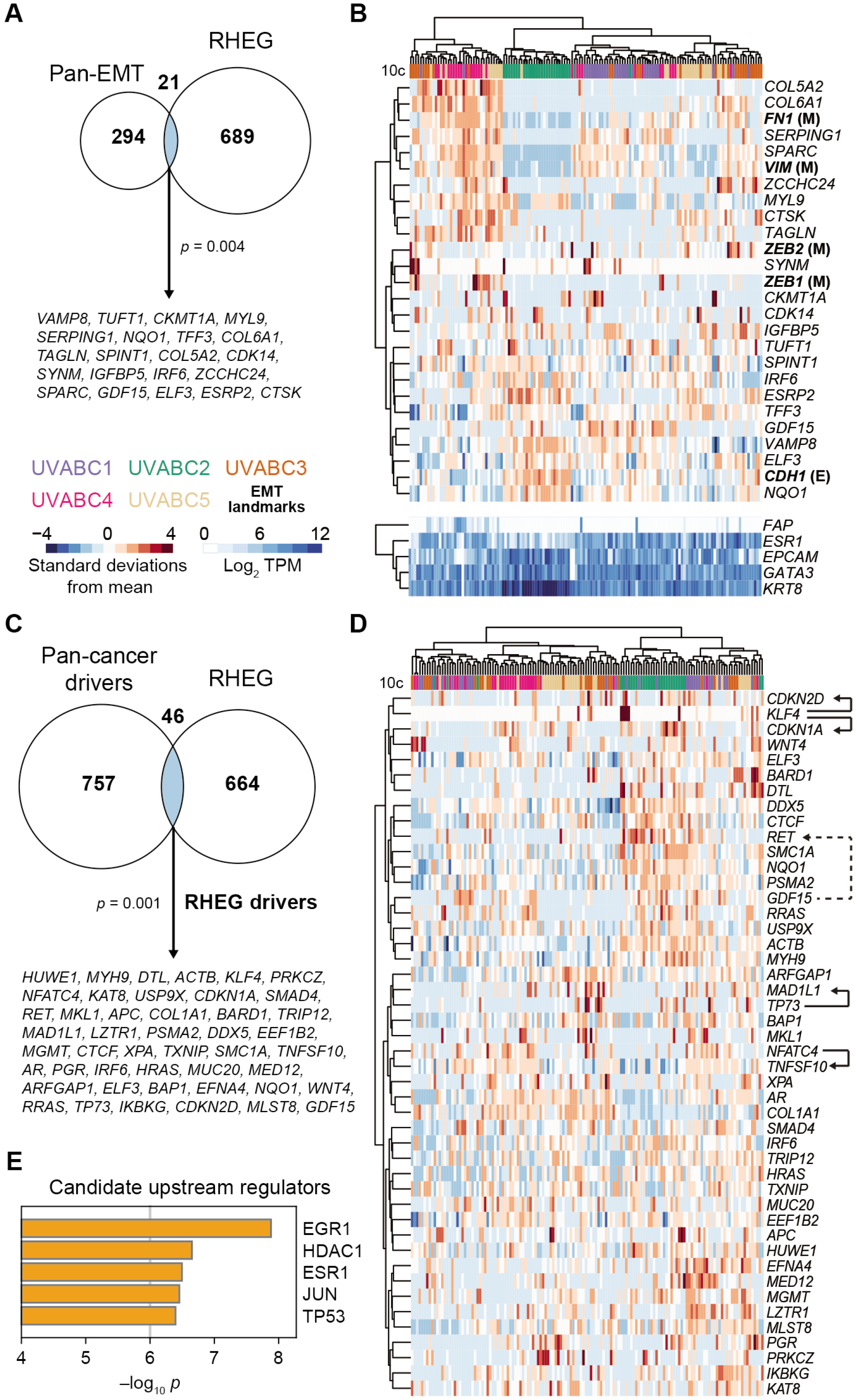
RHEGs contain epithelial-to-mesenchymal transition (EMT) markers and driver genes for cancers other than breast. **A,** Venn diagram intersecting the UVABC RHEG set with a pancancer EMT signature (34). Shared genes are listed. **B,** Hierarchical clustering of the shared genes in **A** along with epithelial (E: *CDH1)* and mesenchymal (M: *VIM, FN1, ZEB1, ZEB2)* markers that were reliably detected by 10cRNA-seq. Stromal character is indicated by the relative abundance of the fibroblast marker *FAP* compared to the luminal markers *ESR1, EPCAM, GATA3,* and *KRT8.* **C,** Venn diagram intersecting the UVABC RHEG set with a pan-cancer set of driver genes (22,35). The intersection was updated to include proximal RHEGs as described in Supplementary Table ST3. Shared genes (“RHEG drivers”) are listed. **D,** Hierarchical clustering of RHEG drivers. Arrows with solid lines between co-clustering drivers indicate possibly direct transcription factor-target gene relationships as described in the text. The arrow with a dashed line indicates a ligand-receptor interaction. **E,** Leading upstream candidate regulators of the RHEG drivers. The full list of candidates is available in the complete Metascape analysis (Supplementary File S9). For **A** and **C**, statistical significance of overlaps was assessed by the hypergeometric test.

### RHEGs are enriched for pan-cancer driver genes and suggest transcription factor-target relationships in single cells

The dearth of breast-cancer driver genes among RHEGs (**Fig. 5B**) prompted us to look at cancer drivers more broadly. We merged 299 robust drivers for any cancer type (22) with the latest pan-cancer analysis reporting 803 drivers from 2,658 tumors (35) and intersected with the RHEG set. There were multiple instances where RHEGs resided in the same complex, pathway, or gene subfamily as a cancer driver (Supplementary Table ST3). We included these proximal RHEGs and altogether found 46 genes as “RHEG drivers” shared between the two datasets (*p* = 0.001 by hypergeometric test; **Fig. 6C** and Supplementary File S8). Even with the expanded driver set, we found no enrichment for mutated breast-cancer driver genes (*p* = 0.6 by hypergeometric test). In the UVABC cohort, RHEG drivers may be leveraged noncanonically through transcriptional regulation rather than mutation.

Last, we clustered the RHEG drivers to ask whether there were interpretable covariations spanning multiple patients (**Fig. 6D**). Associations among 10-cell samples were tightest for UVABC2 and UVABC4, in line with their separation on the earlier UMAP (**Fig. 4D**). Repeatedly, co-clustering RHEG drivers suggested direct modes of action between transcription factors and target genes (**Fig. 6D**, solid arrows). For example, knockdown of *NFATC4* blocks induction of the neighboring RHEG driver, *TNFSF10* (50), and there is literature that the reprogramming factor *KLF4* is required for full induction of *CDKN1A* (51). Although no functional studies are available for *CDKN2D,* another co-clustering cyclin-dependent kinase inhibitor (**Fig. 6D**, solid arrows), the *CDKN2D* locus is occupied by KLF4 (52) and may warrant further study. Likewise, the *MAD1L1* locus is reportedly among the top 250 binding events in the genome for *TP73* (53)—a RHEG driver absent from all luminal breast cancer cells profiled by scRNA-seq (20). Some of the debate involving *MAD1L1* as a TP53 target gene [reviewed in (54)] could be explained by compensation from TP73 (55). The finding is consistent with the *TP53* mutations identified in UVABC1-3 and bioinformatic prediction of the TP53 family as a candidate upstream regulator of RHEG drivers overall (**Fig. 6E** and Supplementary Files S1 and S2). Likewise, given the hormone-positive luminal subtype, we were encouraged to see ESR1 and the ESR1 effector EGR1 (56) as leading candidates for upstream regulation (**Fig. 6E** and Supplementary File S9). RHEG drivers are variably expressed within luminal breast cancers, and our data suggest that some are variably active.

## Discussion

This work combines 10cRNA-seq (13) and stochastic profiling (17) for disruption-free isolation of cancer cell-regulatory heterogeneities in a clinically practicable way. We targeted LCM isolation to breast-carcinoma cells here by using nuclear cytology or epithelial-targeted antibodies; the approach is also compatible with genetically encoded fluorophores (12,40). For 10cRNA-seq, artifactual cell stress (11) is avoided by LCM (12), and dominant cycling transcripts are mitigated by 10-cell averaging that blurs out sample-to-sample differences in cellcycle phase. But in many respects, 10cRNA-seq of malignant cells shares similarities with scRNA-seq: cases are very different from one another, and samples vary substantially within cases. What differs is overall gene coverage per sample (10cRNA-seq > scRNA-seq), as well as the analytical approach needed to discern single-cell differences. Abundance-dependent overdispersion can identify candidates from 10cRNA-seq, much like the nonparametric distribution tests first deployed for microarray data (16). As with microarrays, we anticipate future developments toward parameterizing the underlying single-cell distributions, which combine to yield 10cRNA-seq observations (19). The 10cRNA-seq-based subtype classifications predicted local differences not obvious from histology, and tools for spatial analysis of biomolecules are rapidly advancing (57). It will be especially intriguing when spatial transcriptomics (58) reaches the resolution and sensitivity of 10 cells.

A concern when initiating this study was that copy-number variations within and among primary tumors would dominate the heterogeneities identified by stochastic profiling. Fortunately, gene candidates were uncoupled from chromosomal gains–losses measured in bulk by whole-exome sequencing and estimated at the 10-cell level by inferCNV. Restricting to early-stage cancers was wise, because tumors with more extensive and variegated genomic aberrations would surely have such variation reflected in transcript abundance. Our analyses suggest that UVABC1—with a 10q loss and partial 11q gain typical for the luminal subtype (36)—was on the cusp of skewing toward genomic heterogeneity. As with recurrent copy-number variations, recurrent early-stage heterogeneities in transcriptional regulation may be characteristic of some cancer subtypes.

RHEGs open the possibility of making more specific claims about intratumor heterogeneity beyond cell stress, cell cycle, cell proteostasis, and cell type (9). A recent scRNA-seq study documented twelve such recurrently heterogeneous programs in a large panel of cell lines (59). Although genes in these programs overlap significantly with RHEGs (*p* < 10^-8^ by hypergeometric test), they only comprise ~9% of the RHEG set overall. Partial EMTs in carcinomas have been documented by scRNA-seq (20,38), which we verify here in earlier-stage tumors without any pre-dissociation (**Table 1**). While there are many ways to elicit EMT-like states, a leading explanation for the UVABC cohort is tissue stiffness given their marked desmoplasia. Notably, one RHEG is the hemidesmosomal integrin *ITGB4,* which acts as a critical sensor for matrix stiffness in breast epithelia (3). *ITGB4* was undetected in every luminal breast cancer cell analyzed by conventional scRNA-seq (20).

An important finding from this work is the overlap between RHEGs and pan-cancer drivers. Unlike genomic alterations, transcriptional changes are reversible, rendering them less risky from the standpoint of early tumor evolution. Induced or repressed genes that productively elicit cancer hallmarks have the subsequent opportunity to be “hard-coded” by mutation, deletion, or amplification. Among RHEG drivers, secreted ligands and receptors are not overly prevalent, but a pair with some coordination is the p53 target gene *GDF15* and its cognate receptor *RET* (60) (**Fig. 6D**, dashed arrow). In an accompanying study on another carcinoma type (12), we suggest that such receptor-ligand pairs could engage as locally varying autocrine-paracrine circuits within a tumor that shape the immune microenvironment. For instance, among the 136 candidates shared by the two cases requiring immuno-LCM because of extensive lymphocytic infiltration, we identified an inhibitory ligand for NK cells (*ADGRG1*), a major histocompatibility class II receptor (*HLA-DRA*), a macrophage stimulatory ligand (*MST1*), and the palmitoyltransferase for PD-L1 (*ZDHHC9).* Natural variation in such carcinoma transcripts could one day be mapped to associating changes in the type and extent of immunecell recruitment (61).

Single-cell cancer biology must trade off coverage, throughput, and handling artifacts to retain the conceptual allure of measuring one cell. The approaches described and implemented here for late-onset, early-stage breast cancer are also a compromise, but one that triangulates differently by using cell pools to reduce handling and improve coverage. We see great potential in using stochastic profiling by 10cRNA-seq to deconstruct the very earliest stages of tumor initiation and premalignancy in engineered systems (40) and in precancerous *in situ* lesions of the breast where the need for treatment is actively debated (62).

## Acknowledgments

We thank Emily Farber and Suna Onengut-Gumuscu at the UVA Center for Public Health Genomics for RNA-seq library preparation and sequencing, Craig Rumpel and Angela Miller at the UVA Biorepository and Tissue Research Facility for LCM maintenance and histology services, UVA Research Computing for high-performance computing access and consulting, Henry Pritchard for assistance with NCBI GEO deposition, and Dylan Schaff, Francine Garrett-Bakelman, Hui Zong, and Cheryl Borgman for critical evaluation of the manuscript. This work was supported by the National Institutes of Health #R01-CA194470 (K.A.J.), the David & Lucile Packard Foundation #2009-34710 (K.A.J.), a Wagner Fellowship (S.S.), the UVA Medical Scientist Training Program (S.S.), and a UVA Cancer Center support grant #P30-CA044579.

