## Supplementary Figures and Tables for "Pan-cancer Drivers are Recurrent Transcriptional Regulatory Heterogeneities in Early-stage Luminal Breast Cancer"

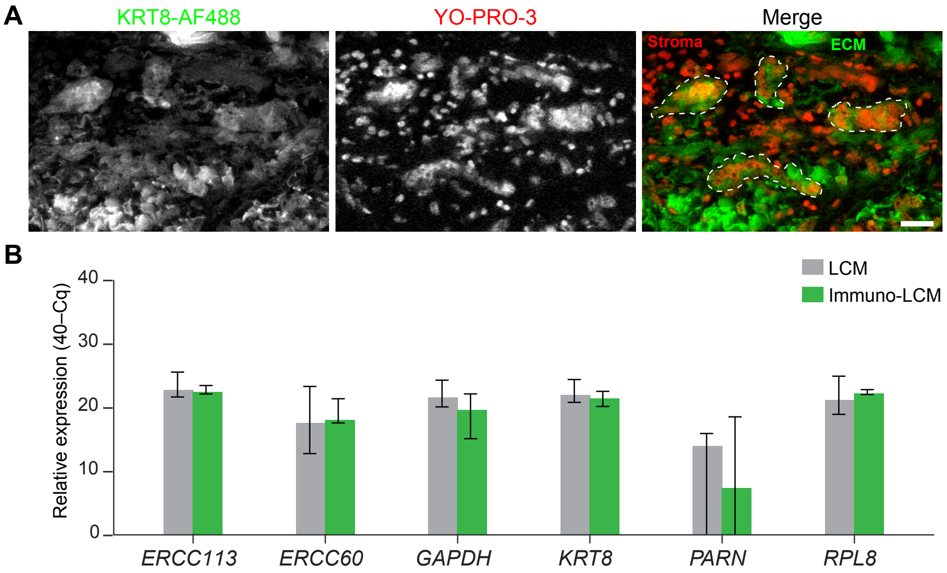


**Supplementary Figure S1.** Immuno-LCM capture of 10-cell samples does not affect gene detection. **A,** Rapid immunostaining of UVABC3 with an Alexa Fluor 488 (AF488)-conjugated antibody recognizing KRT8 (left) combined with YO-PRO-3 to stain all nuclei (middle). Epithelial cells were identified by dual red-green staining (right, dashed lines) compared to stromal cells that are KRT8-negative and autofluorescent extracellular matrix that is free of nuclear staining. Scale bar is 25 µm. **B,** Relative abundance for the indicated transcripts as measured by quantitative PCR in UVABC4. Two exogenously spiked-in RNA transcripts (*ERCC113*, *ERCC60*) were quantified along with four endogenous genes. Data are shown as the median inverse quantification cycle (40–Cq) ± range from $n$ = 4 replicates.

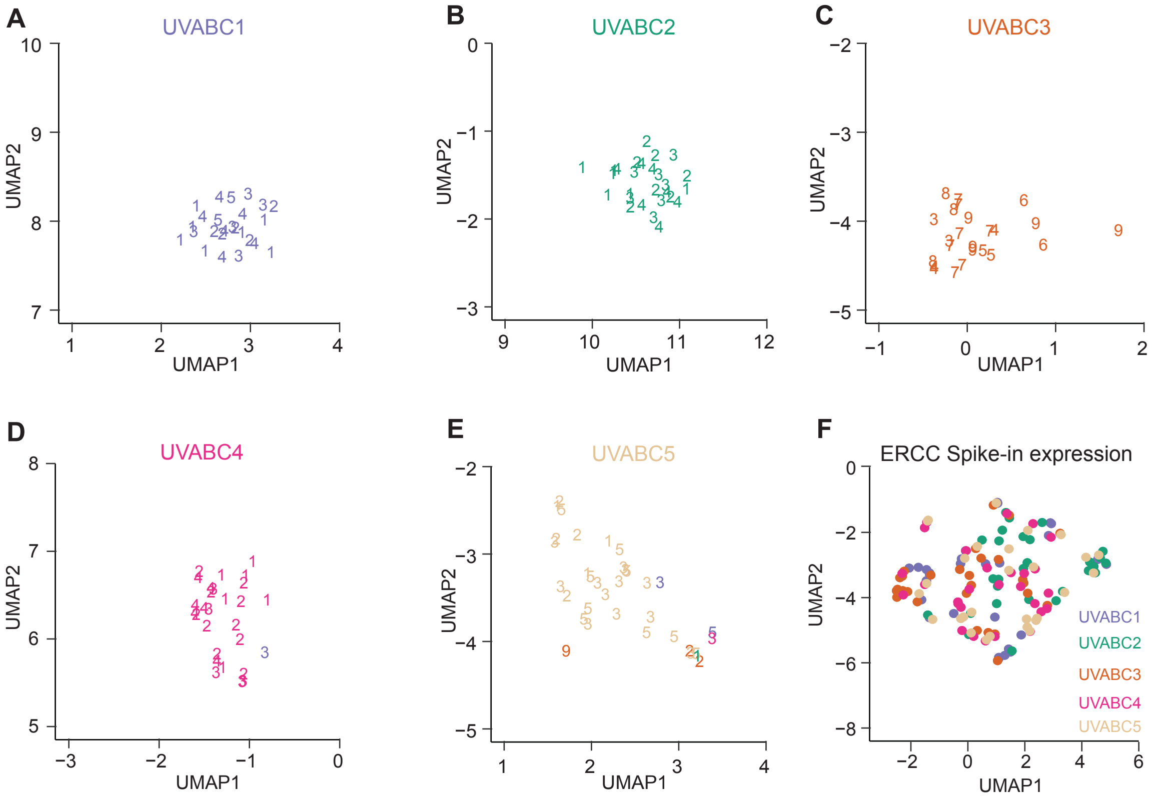


**Supplementary Figure S2.** Clustering of 10cRNA-seq data by tumor does not arise from batch effects. **A–E,** Enlarged UMAP embedding for UVABC tumors from **Fig. 2A**, with different batches of sample collections annotated by number. **F,** UMAP embedding of the UVABC cohort based only on the ERCC spike-in transcripts of 10cRNA-seq samples.

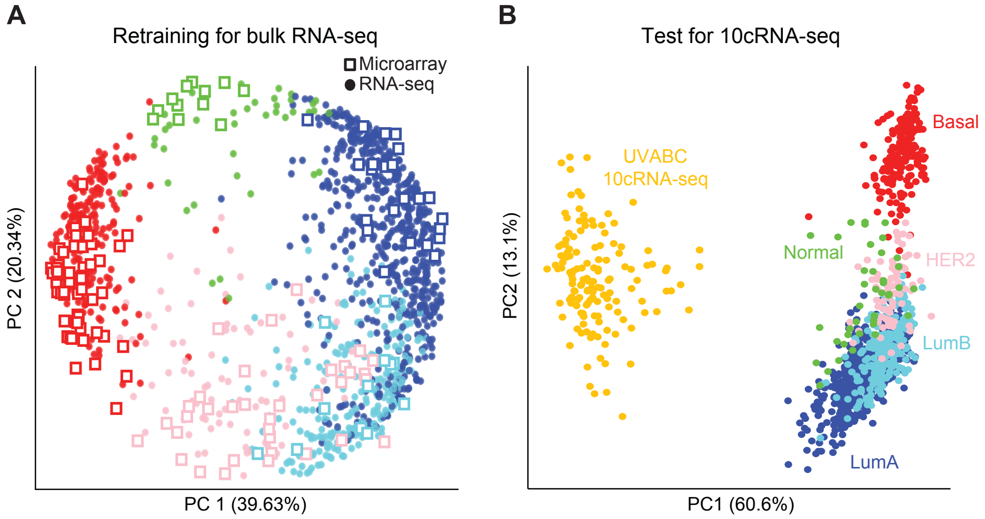


**Supplementary Figure S3.** Microarray-based PAM50 classification is not adaptable to 10cRNA-seq. **A,** Principal component plot adapting the microarray-based PAM50 predictor (1) to bulk RNA-seq data of breast tumors from The Cancer Genome Atlas (TCGA) (2) (see Materials and Methods). Similar projections of the different data types indicate successful data fusion: 77% of RNA-seq samples were correctly subtyped, with an average confidence score of 0.99. **B,** Principal component plot of UVABC 10cRNA-seq observations together with TCGA tumors subtyped with the adapted PAM50 classification of **A**. 10cRNA-seq projections are distinct from all classified subtypes of TCGA tumors, even though dispersion along PC2 indicates underlying subtype differences.

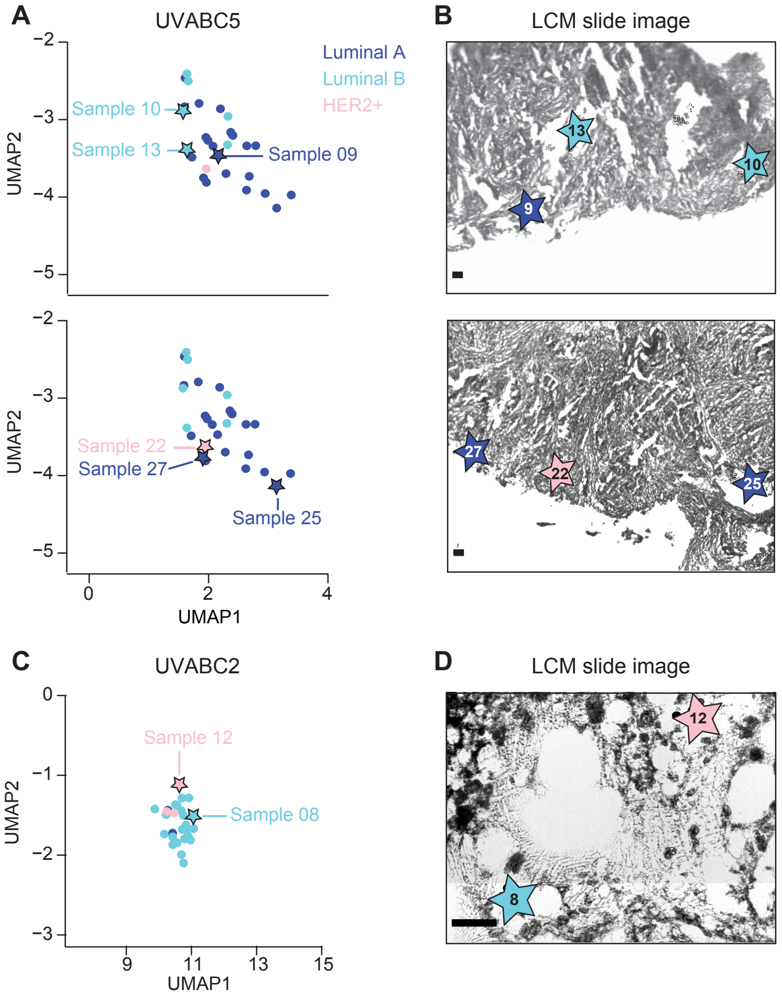


**Supplementary Figure S4.** Different molecular subtypes are assigned to tumor cells microdissected from the same cryosection. **A,** Enlarged UMAP embedding for UVABC5 from **Fig. 2C**, highlighting six 10cRNA-seq samples (stars) obtained together in two separate cryosections. **B,** Low-magnification grayscale LCM slide images indicating the regions microdissected in the UVABC5 cryosections. **C,** Enlarged UMAP embedding for UVABC2 from **Fig. 2C**, highlighting two 10cRNA-seq samples (stars) obtained together in one cryosection. **D,** High-magnification grayscale LCM slide image indicating the regions microdissected in the UVABC2 cryosection. Scale bar is 50 μm (**B** and **D**).

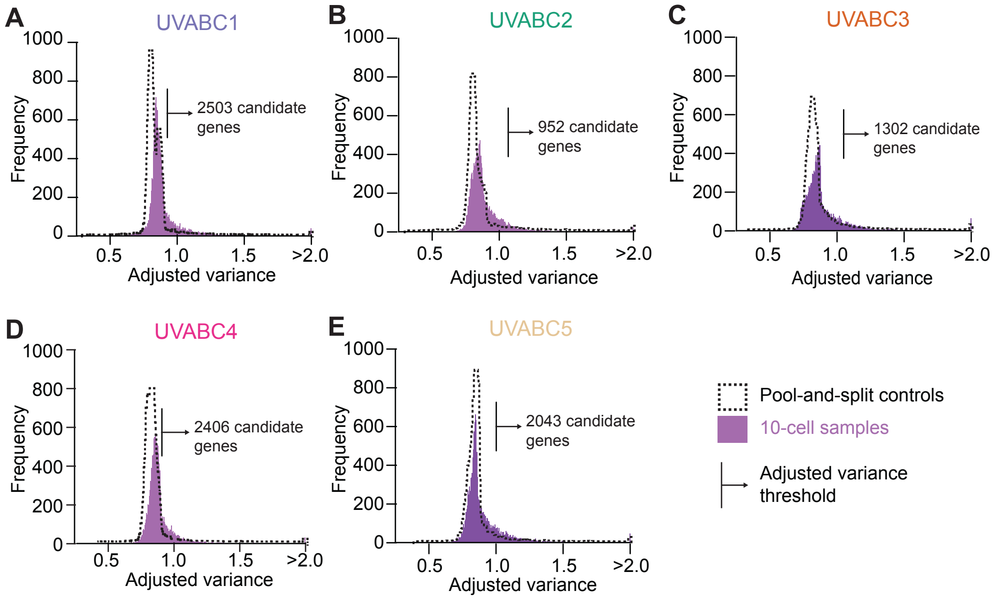


**Supplementary Figure S5.** Abundance-dependent overdispersion statistics of individual cases in the UVABC cohort. **A–E,** Distribution of adjusted variance scores for each gene measured transcriptomically in separate 10-cell samples (purple) compared to pool-and-split controls estimating technical variation (black dashed). The arrow indicates the 5% type I error rate of adjusted variance estimated empirically from the pool-and-split controls, which was used as the cutoff for 10-cell samples.

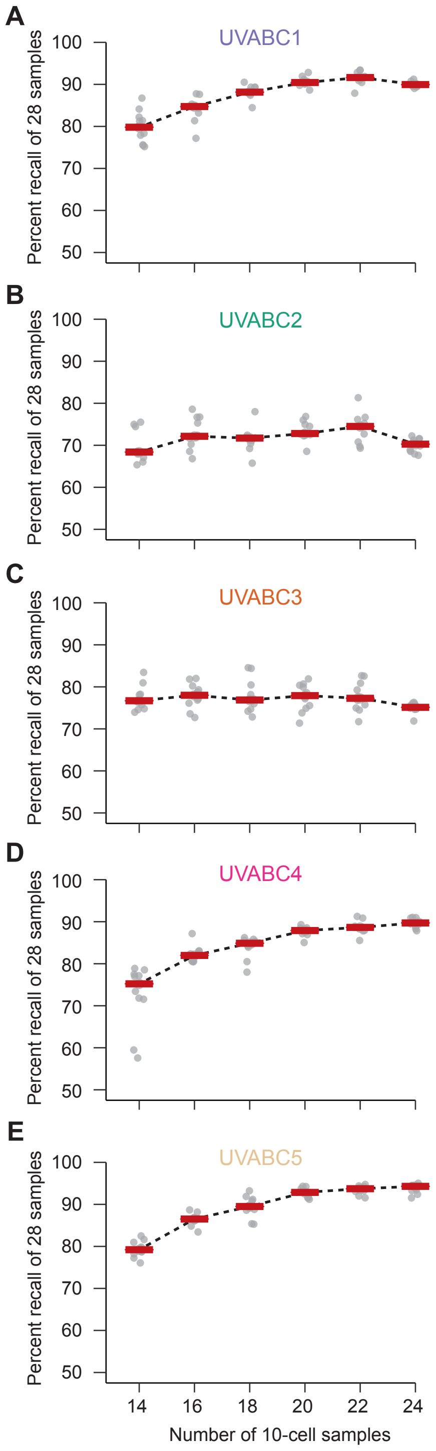


**Supplementary Fig. S6.** Recall of candidate heterogeneities saturates with 20 or greater 10-cell samples. **A–E,** Percentage of candidates for UVABC1–5 recalled as a function of the indicated number of subsampled 10cRNA-seq observations. Results are shown as the median (red) from *n* = 10 computational iterations of subsampling without replacement from the full 28-observation dataset for each UVABC case (jittered markers).

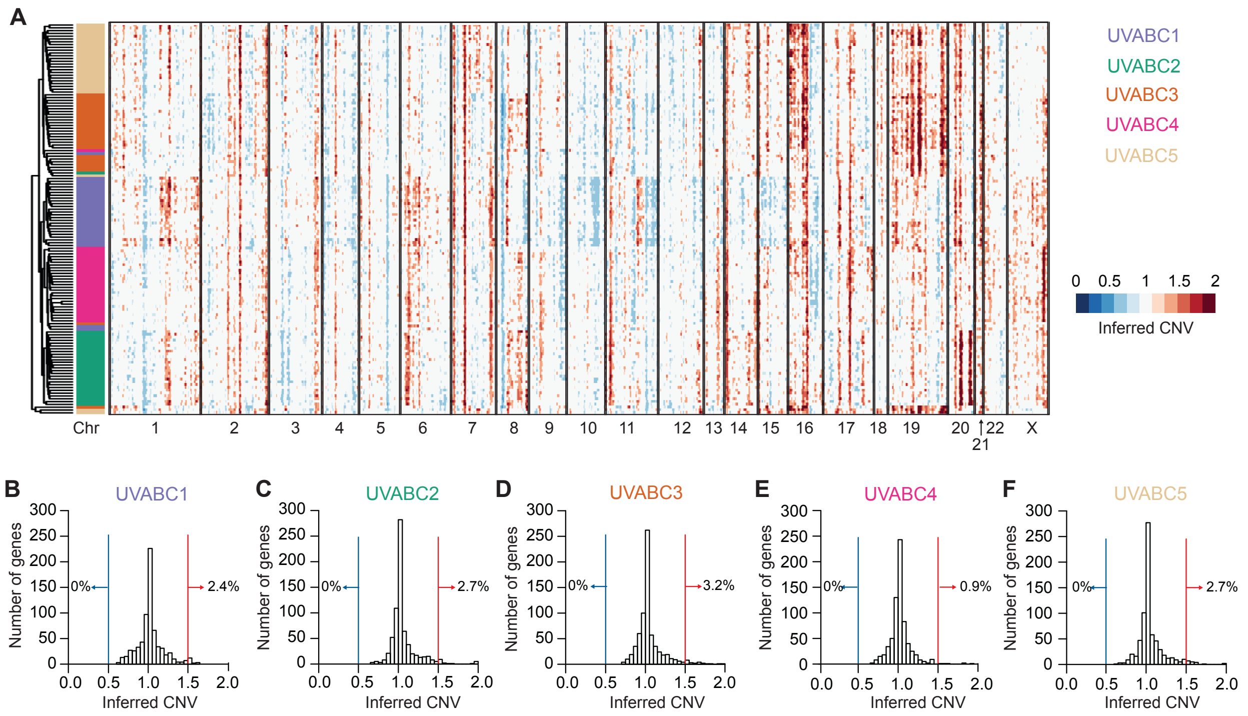


**Supplementary Figure S7.** Most candidate heterogeneities do not reside in loci with inferred copy-number variations (CNVs). **A,** Chromosomal gains and losses predicted from 10cRNA-seq data by inferCNV (3). RNA-seq data from normal human luminal breast tissue obtained through GTEx (4) was used as the reference transcriptome. Gains in 1q (UVABC1, UVABC2) and 8q (UVABC2, UVABC4) and losses in 8p (all) and 16q (UVABC1, UVABC4) are characteristic of luminal A breast tumors (5). **B–F,** Distribution of inferred CNVs corresponding to the candidate heterogeneities of individual cases in the UVABC cohort. Percentage of transcripts with inferCNV scores suggesting gain (red) or loss (blue) is shown.

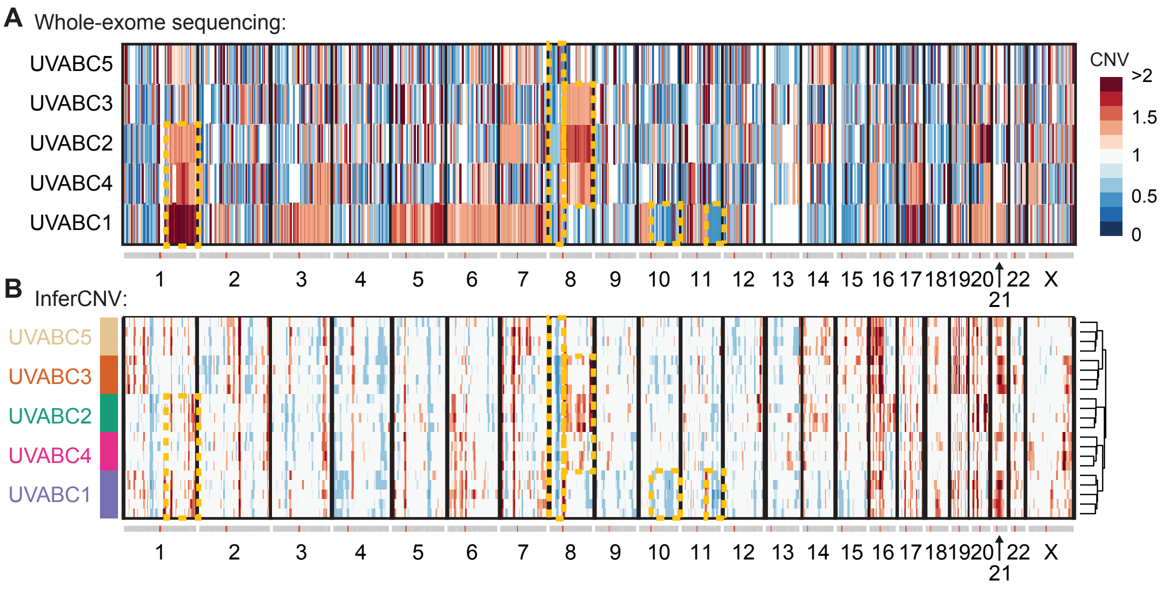


**Supplementary Figure S8.** Whole-exome sequencing confirms copy-number variations (CNVs) predicted by inferCNV. **A,** CNV estimates across the genome from whole-exome sequencing. **B,** CNV estimates from inferCNV (3) using 10cRNA-seq pool-and-split controls ($n$ = 4–5 spatially separated replicates for each UVABC case). For **A** and **B**, chromosomal arms gained or lost by whole-exome sequencing that agree with inferCNV are highlighted in yellow.

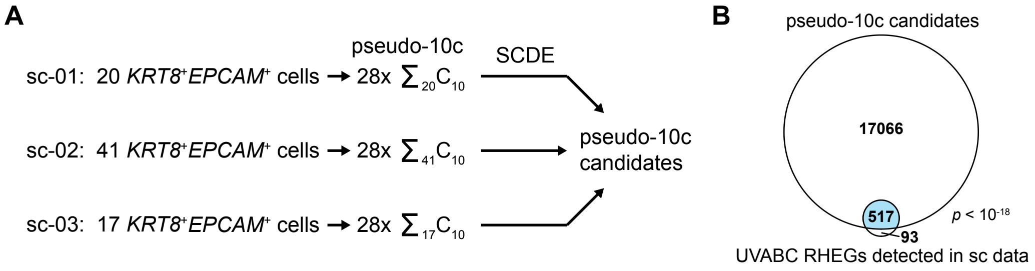


**Supplementary Fig. S9.** RHEGs estimates are stable when extended to non-UVABC cases of luminal breast cancer. **A,** Computational strategy for assembling "pseudo-10-cell" (pseudo-10c) datasets from three cases of luminal breast cancer analyzed by scRNA-seq (6). Only carcinoma cells (defined as cells expressing *KRT8* and *EPCAM* at TPM > 1) were used for pseudo-10c assembly. The same SCDE-based identification of overdispersed transcripts was used with the minimum adjusted variance type I error rate of the UVABC cohort. **B,** Venn diagram intersecting the RHEGs detectable in the pseudo-10c datasets and the pseudo-10c candidates.

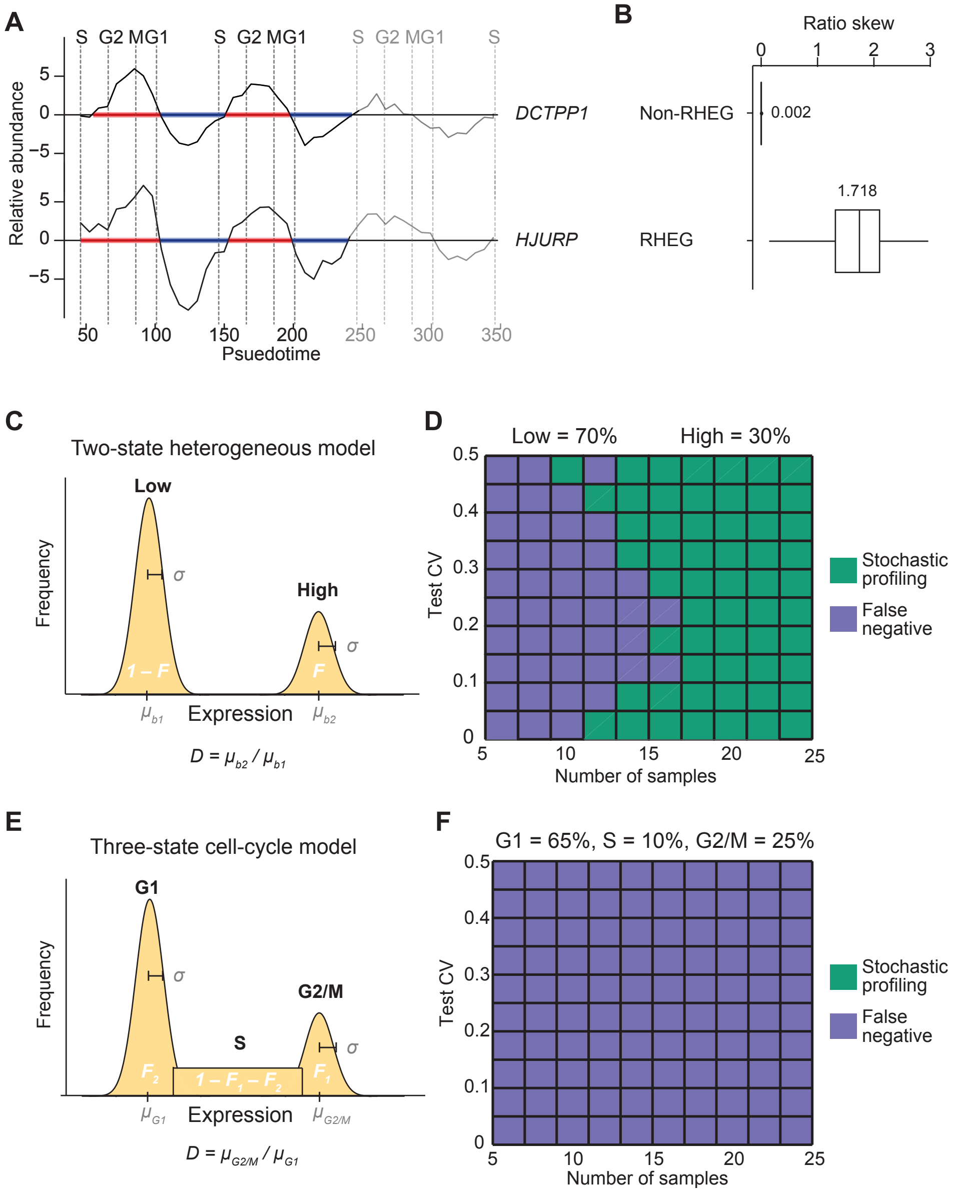


**Supplementary Figure S10.** Periodically cycling transcripts are disfavored by stochastic profiling. **A,** Asymmetric cell-cycle oscillations in the RHEG *DCTPP1* (upper) compared to the non-RHEG *HJURP* (lower). Microarray data from synchronized HeLa cells (7) and pseudotime estimates were obtained from Cyclebase 3.0 (8). Time intervals above (red) and below (blue) the midpoint of each transcript are shown. **B,** Asymmetry of cycling RHEGs quantified by ratio skew of above–below time intervals compared to non-RHEG cycling transcripts (see Materials and Methods). Data are shown as boxplots from $n$ = 1000 bootstrapping runs. **C,** Abstraction of a two-state regulatory heterogeneity as a probability distribution with four parameters: two population means ($\mu_{b1}$ and $\mu_{b2}$, yielding a fold difference $D$), a common log coefficient of variation (= $\sqrt{e^{\sigma^2} - 1}$), and an expression fraction ($F$) capturing the relative proportion of the two regulatory states. Additional details of the probability distribution are described elsewhere (9-11). **D,** Monte-Carlo simulations (10) of stochastic profiling in the two-state case. False-negative regimes are marked when a two-state heterogeneity is not detected in the 10-cell pool. **E,** Abstraction of a three-state cell-cycle model. A uniform S-phase interval is added in between the first and second regulatory states modeling G1 and G2/M phases, and the fractional expression parameter is expanded to $F_1$ and $F_2$ accordingly (see Materials and Methods). **F,** Monte-Carlo simulation of stochastic profiling in the three-state case. The false-negative regime marks three-state heterogeneities that are not detected in the 10-cell pool. For **D** and **F**, the following simulation parameters were used: $D$ = 3, $\sigma$ = 0.2, $F$ = 0.3 (**D**) or $F_1$ = 0.25 and $F_2$ = 0.65 (**F**). Code for the three-state cell-cycle model is available in Supplementary File S5.

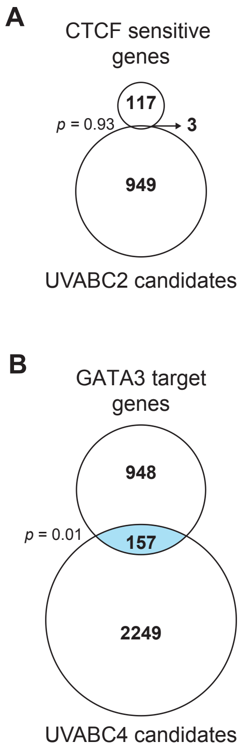


**Supplementary Figure S11.** Intersection analysis of RHEGs with UVABC cases harboring mutant transcriptional drivers for breast cancer. **A,** Venn diagram intersecting the candidate heterogeneities of UVABC2 (harboring a CTCF F416L mutation) with a list of transcripts altered by CTCF knockout in a luminal breast cancer cell line (12). **B,** Venn diagram intersecting the candidate heterogeneities of UVABC4 (harboring a GATA3 D48Y mutation) with a list of GATA3 target genes (13).

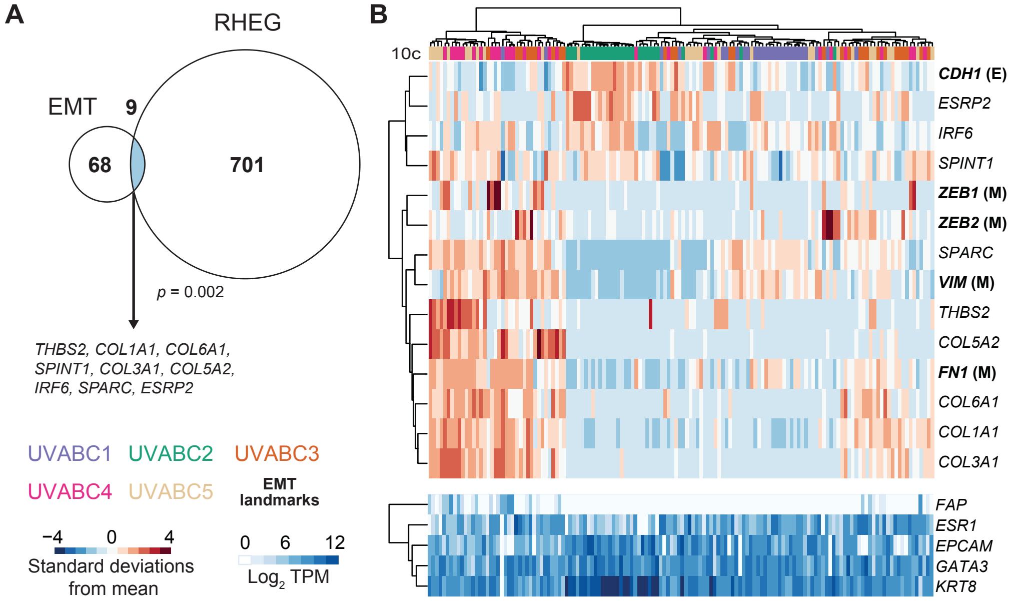


**Supplementary Figure S12.** RHEGs are enriched for additional epithelial-to-mesenchymal transition (EMT) signatures. **A,** Venn diagram intersecting the UVABC RHEG set with an independently derived EMT signature (14) from that of **Fig. 6A**. Shared genes are listed. **B,** Hierarchical clustering of the shared genes in **A** along with epithelial (E: *CDH1*) and mesenchymal (M: *VIM*, *FN1*, *ZEB1*, *ZEB2*) markers that were reliably detected by 10cRNA-seq. Stromal character is indicated by the relative abundance of the fibroblast marker *FAP* compared to the luminal markers *ESR1*, *EPCAM*, *GATA3*, and *KRT8*.

**Supplementary Table ST1.** Early-stage luminal tumors profiled in this study.

| Sample | Age | Race | Stage | Grade | ER | PR | HER2 | Mitoses (per hpf) | Tumor infiltrating lymphocytes foci (per hpf) | Desmoplastic reaction |
|---|---|---|---|---|---|---|---|---|---|---|
| UVABC1 | 76 | Caucasian | 1 | 3 | + | + | — | 21/10 | 2/5 | Marked |
| UVABC2 | 52 | Caucasian | 2 | 3 | + | + | — | 32/10 | 2/5 | Moderate |
| UVABC3 | 63 | Caucasian | 1 | 3 | + | + | — | 11/10 | 5/5 | Marked |
| UVABC4 | 63 | Caucasian | 1 | 2 | + | + | — | 2/10 | 1/5 | Marked |
| UVABC5 | 59 | Caucasian | 1 | 1 | + | + | — | 2/10 | 3/5 | Marked |

hpf = high powered field (400x magnification)

**Supplementary Table ST2.** Candidate heterogeneities are enriched in upregulated transcripts.

| Sample | Heterogeneous | Upregulated | Downregulated | Heterogeneous + upregulated | Heterogeneous + downregulated |
|---|---|---|---|---|---|
| UVABC1 | 2503 | 2912 | 3000 | 674 ($p \ll 10^{-16}$) | 176 ($p \sim 1$) |
| UVABC2 | 952 | 2511 | 3478 | 296 ($p \ll 10^{-16}$) | 29 ($p \sim 1$) |
| UVABC3 | 1302 | 2964 | 4360 | 263 ($p < 10^{-14}$) | 165 ($p \sim 1$) |
| UVABC4 | 2406 | 2857 | 3327 | 555 ($p \ll 10^{-16}$) | 235 ($p \sim 1$) |
| UVABC5 | 2043 | 3515 | 4361 | 487 ($p \ll 10^{-16}$) | 362 ($p = 0.26$) |

**Supplementary Table ST3.** Multiple RHEGs are proximal to established cancer driver genes.

| RHEG | Driver gene | Cancer type(s) | Proximal relationship |
|---|---|---|---|
| *EFNA4* | *EPHA4* | LUAD | EPHA4 signals through EFNA4 |
| *GDF15* | *RET* | PANCAN, THCA, SKCM | RET is a coreceptor for GDF15 |
| *NQO1* | *NFE2L2* | PANCAN, LUSC, LIHC | NFE2L2 activity is marked by *NQO1* abundance |
| *WNT4* | *WNT5A* | PRAD | WNT5A and WNT4 are both non-canonical Wnt ligands |
| *RRAS* | *KRAS* | PANCAN, COAD, LUAD, PAAD, UCEC, ESCA | KRAS and RRAS are both in the Ras family |
| *TP73* | *TP53* | PANCAN, BLCA, BRCA, GBM, COAD, ESCA, HNSC, KIRC, LIHC, LUAD, LUSC, DLBC, OV, PAAD, PRAD, SKCM, STAD, UCEC, SARC | TP53 and TP73 are in the same family of transcription factors |
| *IKBKG* | *IKBKB* | DLBC | IKBKB and IKBKG are in the same IKK complex |
| *CDKN2D* | *CDKN2A* | PANCAN, HNSC, ESCA, LUSC, PAAD, SKCM | CDKN2A and CDKN2D are related CDK inhibitors |
| *MLST8* | *MTOR* | KIRC | MTOR and MLST8 are in the same MTORC complex |

**Supplementary File S1.** Summary of somatic mutations organized by mutation type, gene, and UVABC case. Luminal breast cancer driver genes (15) are highlighted.

**Supplementary File S2.** Summary of somatic mutations organized by somatic variant and UVABC case. For each variant, the classification is reported along with the HGVSp substitution, and predicted effect by SIFT and PolyPhen where appropriate. Read counts for the alternate allele (t_alt_count) are included with the total read count (t_depth) that together estimate the variant allele frequency (VAF). Luminal breast cancer driver genes (15) are highlighted.

**Supplementary File S3.** Summary of candidate heterogeneities identified from stochastic profiling by 10cRNA-seq and organized by UVABC case. RHEGs recurring in at-least three UVABC cases are assembled (Sheet 6) along with RHEGs not detected by scRNA-seq (Sheet 7). For each candidate, the $\log_2$ fold change (log2FoldChange) and adjusted *p* value (padj) are reported when the candidate was reliably detected in the aggregated scRNA-seq data of breast epithelia from three reduction mammoplasties (16).

**Supplementary File S4.** Summary of differentially expressed genes based on a comparison of pool-and-split controls for each UVABC case compared to aggregated scRNA-seq data of breast epithelia from three reduction mammoplasties (16). Shared transcripts that were predicted to be differentially abundant in all five UVABC cases are assembled in the last worksheet. Each transcript reports the $\log_2$ fold change (log2FoldChange) and adjusted *p* value (padj).

**Supplementary File S5.** MATLAB code (saved as a txt file) for the three-state Monte Carlo simulations of stochastic profiling described in Supplementary Fig. S10.

**Supplementary File S6.** Pseudotime estimates for RHEGs classified as cyclers and for ten non-RHEGs classified as top cyclers by Cyclebase 3.0 (8). The IMAGE clone used for the microarray study of synchronized HeLa cells is listed, and data from the first two cycles used for the analysis in Supplementary Fig. S10B are highlighted.

**Supplementary File S7.** Full METASCAPE (17) analysis of RHEGs organized by gene annotation (Sheet 1), categorical enrichments (Sheet 2), TRRUST candidate regulators (Sheet 3), and predicted enrichments for transcription factor (TF) targets (Sheet 4).

**Supplementary File S8.** Summary of proximal RHEG drivers. For each candidate, the $\log_2$ fold change (log2FoldChange) and adjusted *p* value (padj) are reported when the candidate was reliably detected in the aggregated scRNA-seq data of breast epithelia from three reduction mammoplasties (16).

**Supplementary File S9.** Full METASCAPE (17) analysis of proximal RHEG drivers organized by gene annotation (Sheet 1), categorical enrichments (Sheet 2), TRRUST candidate regulators (Sheet 3), and predicted enrichments for transcription factor (TF) targets (Sheet 4).
